# Methylsulfonylmethane: A Potential Dietary Supplement targeting sphingosine kinase 1 involved in Glioblastomamultiforme

**DOI:** 10.1101/2024.09.05.611570

**Authors:** Faizan Ahmad, Anik Karan, Richard L Jayaraj

**Affiliations:** Department of Medical Elementology and Toxicology, Jamia Hamdard University, Delhi, India; Department of Mechanical and Bioengineering, University of Kansas, Lawrence, KS, USA; Department of Paediatrics, College of Medicine and Health Sciences, United Arab Emirates University, Al Ain, UAE; Department of Biochemistry, College of Science, King Saud University, Riyadh 11451, Saudi Arabia

**Author notes:** **Corresponding Author** Faizan Ahmad Department of Medical Elementology and Toxicology Jamia Hamdard University, Delhi India. Equal Contributing Authors (Faizan Ahmad and Anik Karan).

**Keywords:** SphK1, Methylsulfonylmethane (MSM), Glioblastoma multiforme, Dietary supplement, Therapeutics

## Abstract

Methylsulfonylmethane (MSM) is a common dietary supplement mainly used for inflammatory disorders as well as MSM had shown anti-tumor effects on different types of cancers. However, the glioma cell line has not been tested against MSM, and we are reporting it in our study for the first time. This research used an in silico study in which sphingosine kinase 1(SphK1) is used as a therapeutic target which is associated with Glioblastoma multiforme(GBM) SphK1 is pivotal enzyme for sphingolipid metabolism whose high expression level is thought to be associated with cancer alongside other inflammatory diseases and it is a potential drug target for various types of cancer.First, in silico analysis was executed to evaluate the inhibitory effect of MSM on SphK1.Then we further observed the anti-tumor activities of MSM on the C6 glioma cell line. During in silico investigation at the initial stage, we performed molecular docking with Auto Dock Vina followed by molecular dynamics simulation at 100ns with Gromacs Software Package.MSM binds with SphK1 with a docked score of -2.1 kcal mol^1^. During molecular dynamics simulation complex maintain stability at 10ns but we ran simulation till 100ns to confirm the stability. We performed in depth analysis which includes post trajectory analysis like free energy landscape (FEL), principal constant analysis (PCA) with kernel density (KDE)estimation plots as well as probability distribution plots. Even molecular dynamics simulation shows stability, compactness and interaction of MSM with Sphk1, we calculated MMPBSA binding energy calculation is -13.922 +/- 19.518 kJ/mol^-^ The viability and cellular metabolic activity of the C6 glioma in the presence of MSM showed 393.459 mM/ml of MSM reduced cell viability by 50% (CTC_50_) value in dose dependent manner. Further analysis like DNA fragmentation assay and Acridine orange and ethidium bromide (AO/EB) staining were carried out, which clearly depicts MSM inducing apoptosis in C6 gliomas. Based on in silico and in vitro results,for the first time we are reporting it in our study and we reach to conclusion that that MSM acts as a potential inhibitor for SphK1 as well as inhibits the growth of glioma cells and acts as a potential dietary supplement for the management of GBM which can cross blood brain barrier (BBB) and not toxic to cells even at high doze.

## 1. Introduction

Noticeably prevalent and lethal central nervous system tumours in a grown person is glioblastoma multiforme (GBM), which occurs annually in about 1 in 33000 persons [1, 2]. GBM is extremely deadly, with a median survival of 10-12 months, as low as 6%. GBM is treated by surgical resection, chemotherapy, or both [3]. Moreover, no effective drug is currently available to treat GBM, so we must do thorough research to develop it and bring it into the market. In the current study, we took Methylsulfonylmethane (MSM), a naturally occurring organosulfur compound used as a dietary supplement. MSM has multifaceted biological activities, including anti-tumour and anti-inflammatory activities that relieve stress, increase energy, enhance circulation and metabolism, and effectively heal wounds [4]. Based on the previous study, it has been seen that MSM shows a positive response towards arthritis and multiple inflammatory disorders like intestinal cystitis, allergic rhinitis, and exercise-used inflammation, as well as effective in various cancers like breast cancer [4]. Based on previous findings, MSM is considered a novel compound, and sphingosine kinase1(Spkh1) is a protein of interest for further studies. SphK1 is synthesised as a polypeptide consisting of 384 amino acid residues. It doesn’t possess a transmembrane domain, but it does contain three sequences capable of binding calcium/calmodulin, along with a diverse array of phosphorylation sites for kinases [5]. It is widely expressed in the kidney, liver, lungs, heart, and spleen. Various external stimuli, including the proinflammatory cytokine, tumour necrosis factor, and epidermal and nerve growth factors, activate SphK1[5]. SphK1, typically present in the cytosol, is translocated to the plasma membrane when activated. Spkh1 controls tumour cell migration and invasion in a range of haematological and solid malignancies by acting as an oncogene, as Spkh1 is linked to inflammatory diseases, neurodegenerative disorders, diabetes and pulmonary hypertension [5]. Spkh1 is widely known for developing anti-cancer drugs, and it is a protein target with both N and C Terminal Domain with ATP binding sites present in the cavity of N and C domains. Numerous cellular kinases like PKA, CIB1 (calcium- and integrity-binding protein 1), and PKC activates Spkh1[6]. Regulation of inflammatory pathways could promote the progression of GBM, too [6]. ERK1/2 enhances SphK1 activity through the phosphorylation of Ser225, functioning as an oncogene pivotal in regulating cell migration and invasion, particularly in solid tumours [6]. Reactive oxygen species stimulate SphK1 under hypoxic conditions [7], while suppressing SphK1 expression has been shown to hinder hepatic tumorigenesis in diethyl nitrosamine-treated mice [8]. Apart from SKI-II and CB5468139, which compete at the binding region of ATP, the majority of SphK1 inhibitors consist of sphingosine cognates or non-lipid tiny molecule inhibitors that occupy the “J-shaped” substrate-binding region of SphK1[6]. Deeba et al. in 2020 have shown two bioactive ZINC05434006 and ZINC04260971 natural compounds act as a potential Spkh 1 inhibitors, which may lead to the development of effective therapies for GBM after rigorous successful completion of preclinical and clinical studies [9]. Another study by Sonam et al. in 2022 showed that cinchonine and colcemid act as potent drug molecules for the treatment of cancer, which binds to the active site of SphK1, which leads to conformational changes [10]. FTY720, a structural analogue of sphingosine, augments the responsiveness of prostate cancer cells to radiotherapy by acting as an agonist for S1PR1. Furthermore, research has demonstrated that SK1-I, a selective inhibitor of SphK1, hinders the growth of GBM cell lines and suppresses vascularization and tumours development in a mouse model [11]. Despite the pivotal walk-on part of SphK1 in cancer biology, only a few compounds have progressed to clinical trials, mainly because there are some deficiencies in the necessary welfare profile, oral accessibility, and effectiveness in most of the inhibitors in animal models [12]. Numerous essential substances and their by-products recently target the SphK1/S1P pathway [9]. Compared to traditional chemotherapy, they have been demonstrated to be more effective, have fewer adverse effects, and have improved bioavailability [13, 14]. The immense importance of essential substances encouraged researchers to investigate them to find SphK1 inhibitors that are effective in SPHK1-aided GBM cell growth and metastasis. Elevated expression of SPHK1 led to the activation of the PTX3 gene, known as a biomarker for inflammatory diseases responsible for regulating inflammation and complement activation. Notably, SPHK1 and PTX3 worked in tandem, establishing a positive feedback loop that amplified the expression of both proteins within GBM cells. This suggests that targeting SPHK1 could hold promise as a therapeutic approach for treating GBM. In a recent study Miranda et al., 2021, it was observed that the effect of small molecule CBL0137 increases the level of DNA damage and efficacy of radiotherapy for glioblastoma [15]. In another study, Rebekah et al., 2014, developed a small molecule G6 and G6 that showed promising results in T98G cells by reducing the phosphorylation of Jak2 and STAT3 in a dose-dependent manner [16]. In this study, we are focusing on the MSM as a dietary supplement which can be an easier approach to control the adverse progression of gliomas. In current study, we are using MSM because of its uniqueness and there is a limitation of small molecule which is effective on GBM. MSM is a small molecule as well as dietary supplement which is not toxic even at high doses. In current study, we are using MSM which is a dietary supplement, and it show promising result which is mostly used to get control over inflammation, as well as it, has the potential to cross blood brain barrier (BBB). However, MSM anticancer activity and its mode of action have never been studied in relation to C6 glioma cell line. To support our study, we first investigated MSM binding mechanism and inhibitory efficacy towards Spkh1. To observe mechanism and efficacy first, we performed molecular docking to evaluate the binding affinities of MSM against SphK1, followed by molecular dynamics simulation for 100ns using Gromacs and complex maintain stability at 10ns.To look over stability, compactness and interaction we performed in depth analysis of our complex which we ran till 100ns instead of running longer trajectory which includes post trajectory analysis like free energy landscape (FEL), principal constant analysis (PCA) with Kernel density estimation (KDE) plot as well as probability distribution plots. Further, we performed in-vitro studies to observe the effect of MSM on the C6 glioma cell line to validate our in-silico findings. To determine the MSM potential for cytotoxicity, MTT assay was conducted. MSM was found to have an inhibitory concentration (IC50) of 393.459 mM for C6 glioma cells. Following the examination of the cytotoxicity, the apoptotic characteristics were examined using DNA fragmentation ladder assay as DNA fragmentation into oligonucleosomal ladders is characteristic of programmed cell death and to support our findings further, we performed Dichlorodihydrofluorescein diacetate (DCFHDA) staining which depicts elevated reactive oxygen species (ROS) levels in cells as well disruption of the membrane of mitochondria are considered as a hallmark of apoptosis. Finally, we performed Acridine orange and ethidium bromide (AO/EB) staining in which nuclear deformation of cells occur which leads to induced cell death. Lastly, this study proves that MSM can be used for GBM as well as it is not toxic at a high dose for normal cells [4] and effectively binds with Spkh1 making MSM a potential inhibitor. However, validation of MSM is still needed through intense pre-clinical studies before it proceeds to the clinical trials of GBM.

## 2. Materials and Method

### 2.1. ADMET analysis of MSM

Swiss ADME webserver was used to observe MSM pharmacokinetics and drug-likeness properties.

### 2.2. Molecular Docking of MSM against Crystal structure of Sphingosine Kinase 1

Molecular docking using AutoDock Vina was executed on the ligand (MSM) against the 3D structure of Sphingosine Kinase 1 (PDB ID 3VZB). First, the 3D structures of the ligands were obtained from PubChem and prepared for docking using the Open Babel software. The 3D structure of the Sphingosine Kinase 1 receptor was also prepared in Autodocktools. The AutoDock Vina docking protocol was configured, which involved specifying the search space and determining the grid box dimensions according to the active site residue ARG185. Subsequently, the docking runs were started. After conducting the docking, the obtained poses were scrutinized for their binding affinities, and the most promising ones were chosen for subsequent investigation. Utilizing PyMOL software, the binding orientation and interactions between the ligands and the receptor were thoroughly examined. The outcomes of the docking study played a pivotal role in assessing the ligand’s potential as an inhibitor for the crystal structure of Sphingosine Kinase 1[17].

### 2.3 Molecular Dynamic Simulation at 100 nanoseconds with GROMACS

Molecular dynamics (MD) simulation is a computational technique that uses Newton’s laws of motion to study the movement of atoms in a molecule. In this case, the simulation was performed using the Gromacs software package, which is a widely performed and well-established MD simulation software. The first step in the simulation process was the minimization of the protein-ligand complex in a vacuum. This was done performing the steepest descent algorithm, which involves iteratively adjusting the atomic coordinates of the complex to minimize the system’s potential energy. After the minimization, the complex was dissolved in a periodic box of water using the SPC water model. The SPC water model is a simple model that represents water molecules as a single-point charge and is often used as a starting point for more complex water models. The felicitous amount of sodium chloride is added to maintain the complex at a salt concentration of 0.15 M. The resulting complex was then subjected to an NPT (constant pressure, constant temperature) equilibration phase, followed by a production run for 100 ns (nanoseconds) in the NPT ensemble. The NPT ensemble is used to imitate systems at constant temperature and pressure, which are commonly encountered in biological systems. Finally, the trajectory of the simulation was analyzed using various tools provided by the Gromacs software package, such as the protein root mean square deviation (RMSD), root mean square fluctuation (RMSF), the radius of gyration (RG), solvent accessible surface area (SASA), and hydrogen bonding (H-Bond). These analyses allow researchers to study the simulated system’s various structural and dynamic properties, such as its overall shape, flexibility, and interactions with the surrounding solvent 17-20]

### 2.4. Preparation of MSM

2M MSM (Sigma, USA) was weighed and dissolved in Ham’s F12 medium with 2% inactivated FBS (Sigma, USA) to get 2M/mL stock solution. Further, dilutions were prepared from the stock solution to get lower concentrations for cytotoxicity testing, followed by sterilization using a syringe filter.

### 2.5. Cell Culture of C6 glioma cell line

The C6 cell line (NCCS, Pune, India) was cultured in Ham’s F12 media with 10% FBS (Sigma, USA), penicillin (100 IU/mL), streptomycin (100 g/mL), and amphotericin B (5 g/mL) until confluent. The culture environment included a humidified atmosphere with 5% CO2. Cells were dissociated using a TPVG solution, consisting of 0.02% trypsin, 0.02% EDTA, and 0.05% glucose in PBS. The experiments were carried out in 96-well microtitre plates, while the initial cultures were grown in 25 cm2 culture flasks.

### 2.6. MTT Assay

Trypsinization was performed on the monolayer cell culture, and trypan blue was used to count the cells. 10,000 cells were put into each well of the 96-well microtitre plate.The media in each well is taken out and once rinsed with the medium after being in place for 24 hours. To each well, MSM was added at varying concentrations. For comparison, the untreated cells were kept around as a control. The microtitre plate was then nurtured at 37° C for 24 hours in a 5% CO2 environment. Following a 24-hour period, the media in the plate were removed, and 100 µl of MTT in DPBS was put to every well. The plate is shaken feebly and a 3-hour incubation period at 37° C with a 5% CO2 atmosphere. The plates were gently frenzied while 100 µl of DMSO (Sigma, USA) was put to liquify the formazan crystals. A microplate reader (Biotek, USA) was used to measure absorbance at 570 nm [21].

### 2.7. DNA fragmentation

60 mm tissue culture dishes containing C6 cells (3 x 106 cells/ml) were planted and nurtured at 37° C for 24 hours in a 5% CO2. After MSM treatment, the cells underwent a further 24 hours of incubation as before. After the process of incubation, the chromosomal DNA of the cell was arranged by the Genomic DNA Purification Kit (Qiazen, USA). On a 1.8% agarose gel, total genomic DNA was isolated and resolved. Using ethidium bromide staining and a UV transilluminator, we could observe and capture the photographs of apoptotic DNA fragmentation.

### 2.8. Intracellular ROS

To determine the formation of ROS, a DCF assay (Sigma, USA) was performed. Concisely, C6 cells were planted in a 6-well plate at a concentration of 2 × 105 cells/well in a 2mL cell culture medium with 10% FBS. Then, MSM at concentrations of 400 and 600mM was added to the cells and was kept for 24 hours. The cells were stained with 10μM DCFDA (Sigma, USA) dissolved in DMSO and nurtured for 45 minutes at 37°C. Then, PBS was added to the cells twice to wash. ROS production was visualized and captured using fluorescence microscopy (Leica DMI6000B). The fluorescence intensity was measured using ImageJ 1.50b from NIH for quantification of the ROS levels under experimental conditions.

### 2.9. Mitochondrial membrane potential (MMP)

The potential of the mitochondrial membrane was visualized using Rhodamine 123 staining. C6 cells were planted into a 6-well plate containing 2 × 105 cells in each well and cultured for 24h. Then, MSM at concentrations of 400 and 600mM was added to the cells and was kept for 24 hours. The cells were then washed with 1x DPBS and nurtured with Rhodamine 123 (10μM) (Sigma,USA) for 15 minutes at 37°C. After cleansing off the dye, the cells were portrayed using fluorescence microscopy (Leica DMI6000B). The fluorescence intensity was measured using ImageJ 1.50b from NIH for quantification of the MMP activity under particular experimental conditions.

### 2.10. AO/EB staining

Cells were cultured at 2 × 105 cells/well and incubated with MSM (600mM & 400Mm) for 24h. After the process of incubation, cells were cleansed in DPBS and stained with 1mL each of AO (100µg/mL) and EB (100µg/mL) solution (Sigma, USA) for 5 minutes at a temperature of 37°C. After cleansing off the dye, the cells were portrayed using fluorescence microscopy (Leica DMI6000B, Leica, USA). The fluorescence intensity was measured using ImageJ 1.50b from NIH for quantification of the ROS AO/EB staining of gliomas under experimental conditions.

### 2.11 Stastistical Analysis

Statistical inspection was conducted by Minitab 20 software from Minitab LLC., PA, USA and in Microsoft Excel. The results were staged as mean ± standard error (n=3). Statistical significance was estimated performing RM two-way Anova with Geisoor-Greenhouse correction, Tukey’s multiple comparison tests (applied for n=3), and Brown-Forysthe and Welch Anova (n=1).

## 3. Results and Discussion

### 3.1. ADMET analysis of MSM

The ADMET features of the MSM were assessed by using the SwissADME server. MSM shows high GI absorption and the ability to cross the blood-brain barrier. It follows Lipinki’s rule of five without any violations and does not hamper CYP1A2, CYP2C19, CYP2C9, and CYP3A4. The outcomes of the ADMET inspection are presented in table 1

**Table 1:**
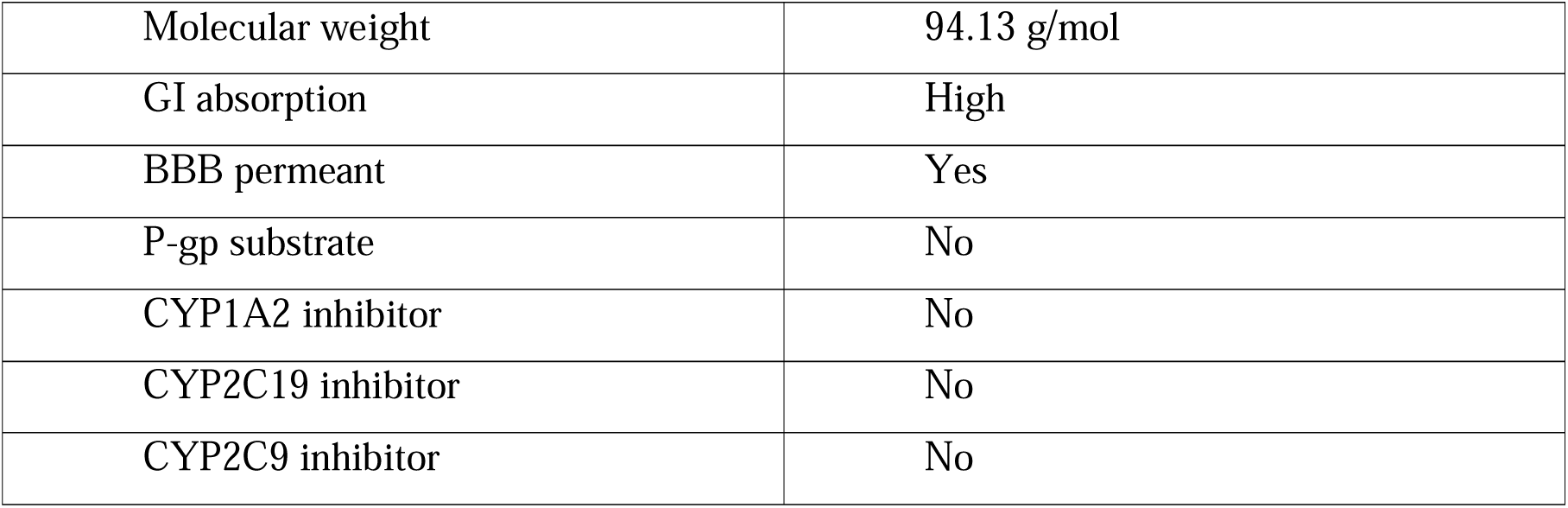
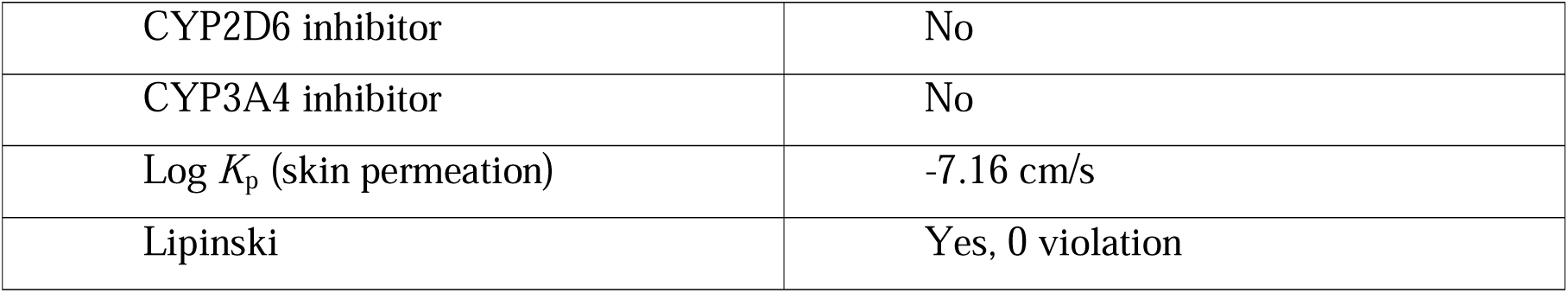
ADMET analysis of MSM with SwissADME.

|  |  |
| --- | --- |
| Molecular weight | 94.13 g/mol |
| GI absorption | High |
| BBB permeant | Yes |
| P-gp substrate | No |
| CYP1A2 inhibitor | No |
| CYP2C19 inhibitor | No |
| CYP2C9 inhibitor | No |
| CYP2D6 inhibitor | No |
| CYP3A4 inhibitor | No |
| Log $K_p$ (skin permeation) | -7.16 cm/s |
| Lipinski | Yes, 0 violation |

### 3.2. Molecular Docking of MSM against Crystal structure of Sphingosine Kinase 1

In this study, ATP and MSM [CID ID-6213] ligand was assessed for their capability to bind to the 3D design of Sphingosine Kinase 1. The compounds’ binding interactions were assessed through a computational method known as molecular docking, which was executed using Auto dock Vina software. Table 2 displays the docking scores of ATP and MSM for the test proteins.

**Table 2:** Docking score and type of interaction of MSM against Crystal structure of Sphingosine Kinase 1.

| Ligand | ID | Affinity kcal/mol | Interaction |
| --- | --- | --- | --- |
| ATP | 5957 | -7.1 | LYS27, SER112, ARG185, ARG191, GLY342, ASN22 |
| MSM | 6213 | -2.1 | ARG185, ARG186 |

The docking results showed that the phytochemicals were similarly effective inhibitors of the Sphingosine Kinase 1 compared to the ATP and MSM. The ATP had docking scores of -7.1 kcal/mol, and MSM had docking scores of -2.1 kcal/mol. Both hydrogen bonding and hydrophobic interactions are observed in between the protein and its ligand. The binding pose, or the specific orientation of the compound with the protein, is depicted in Figure 1.

**Figure 1.**
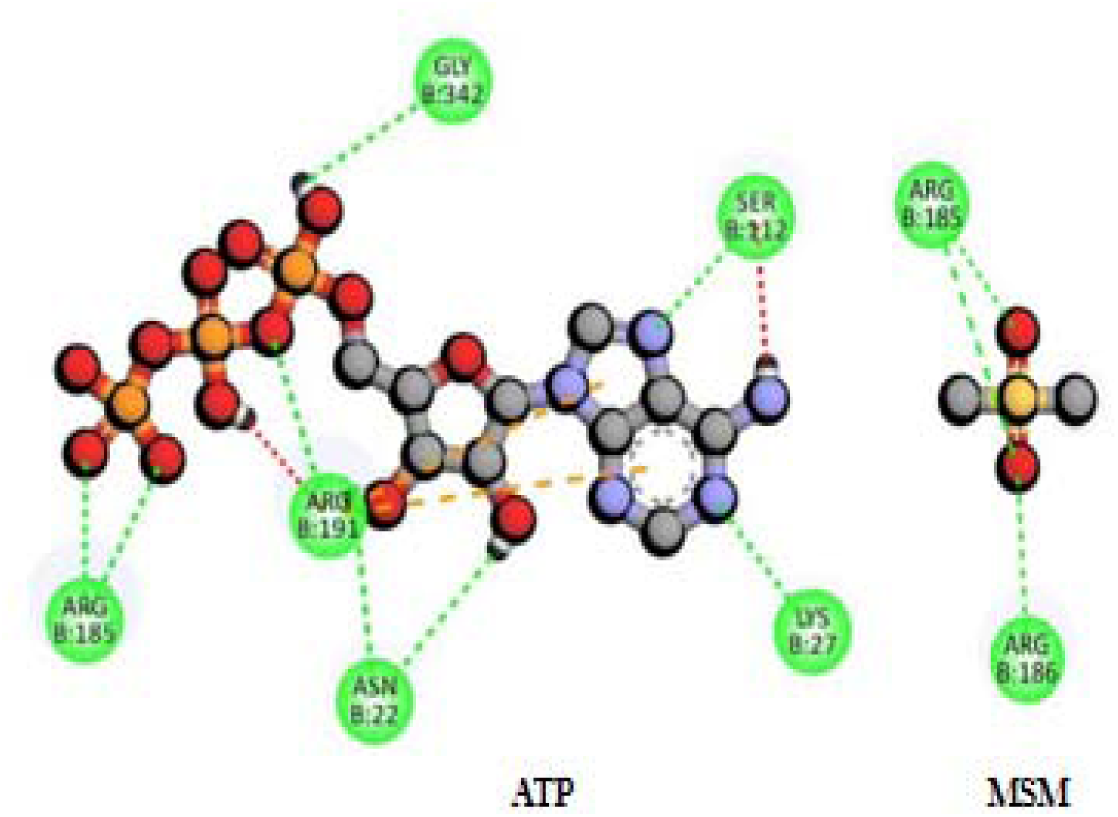
Interaction of ATP and MSM inhibitor at the Sphingosine Kinase 1 active site

### 3.3. Molecular Dynamics Simulation analysis

The use of all-atom MD simulation is a suitable method to examine dynamics of proteins and their interactions with ligands. In this study, MD simulations were employed to examine the dynamic alterations that takes place from the binding of the target protein. Various parameters were computed for both the protein and the protein-ligand complex. Additionally, assessment of the main part inspection and free energy sceneries were studied based on the 100 ns trajectory of the simulation. The RMSD values were monitored over time to inspect the firmness of the SphK1 complex and to observe the system’s behavior, as illustrated in Figure 2A. The findings indicate that both systems reached a state of equilibrium within 10 ns and maintained a consistent distribution throughout the simulation. Moreover, the analysis of RMSD values indicated that the 3VZB-APO, 3VZB-ATP, and 3VZB-MSM complexes retained stability for the entire 100 ns durations, and we did not proceed to longer run, as 100 ns run signifying the enduring firmness of the docked complex. Furthermore, the RMSD pattern of 3VZB-APO demonstrated a reduction after the binding of ATP and MSM in comparison to its free form. This observation suggests that during the simulation the docked complexes of 3VZB-ATP and 3VZB-MSM represent stable systems with minimal fluctuations. The distribution of RMSD values vividly presents that the 3VZB complex exhibited greater stability when bound to ATP and MSM, as depicted in Figure 2.

**Figure 2.**
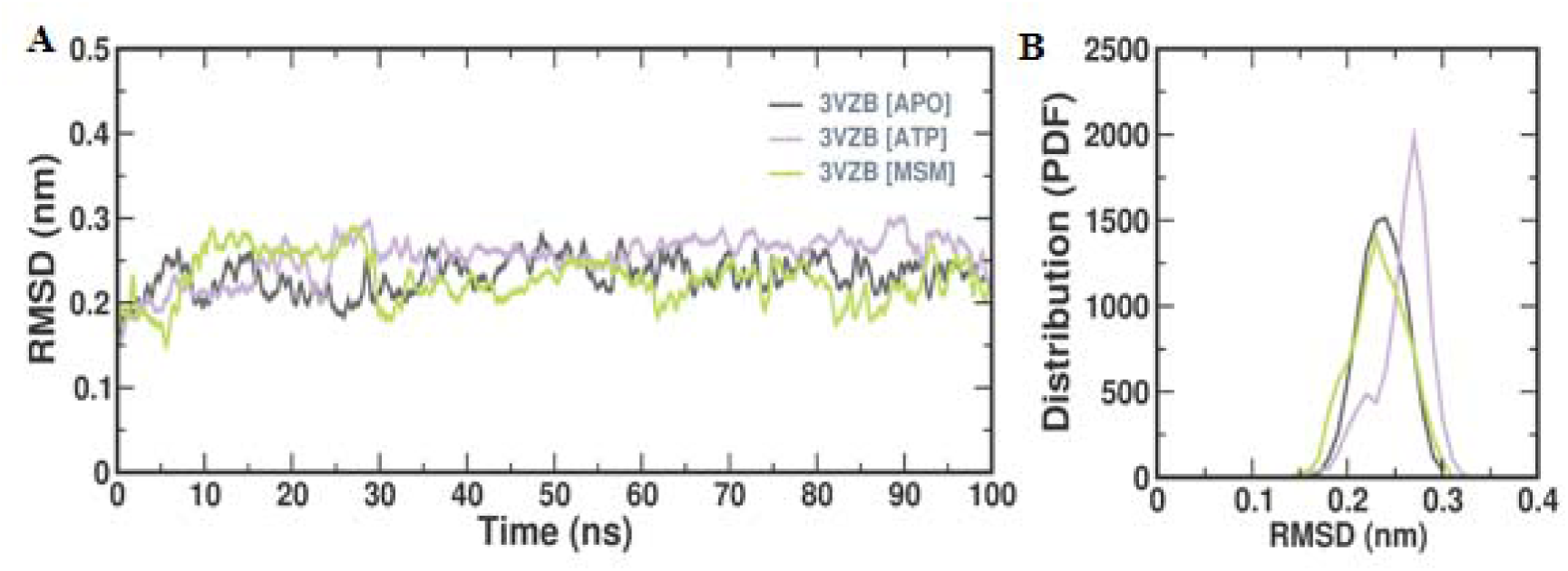
Conformational dynamics of 3VZB-APO, 3VZB-ATP, and 3VZB-MSM complex (A) Time evolution of the RMSD values (B) Probability distribution of RMSD

RMSF is employed to gauge the fluctuations of individual residues and flexible areas within a protein during MD simulations. It enables the assessment of how ligand binding affects the protein. Generally, well-structured elements like sheets and helices exhibit lower RMSF values, while more loosely arranged loop regions tend to have higher RMSF values. In this study, the RMSF values for each compound were determined and plotted for every residue in 3VZB-APO, both beforehand and afterward ATP and MSM binding which is shown in Figure 3. The results indicate that the binding of ATP and MSM did not significantly alter the overall RMSF distribution. Additionally, the ATP and MSM complexes displayed lower fluctuations compared to free APO. To evaluate the dynamic stability and compactness of APO, as well as its ATP and MSM complexes, Rg values were computed and plotted over time, as illustrated in Figures 4A and 4B. The average Rg values for 3VZB-ATP, APO and 3VZB-MSM were determined to be 2.12 nm, 2.15 nm, and 2.13 nm, respectively. The complex system exhibited little reduced Rg values compared to the apo form of 3VZB, indicating a higher level of compactness in the complex. The plot confers that the docked complexes of ATP and MSM with 3VZB maintained stability throughout the entire simulation. Additionally, the probability density function (PDF) plot indicates that APO maintained stability in the distribution of Rg values, with only slight fluctuations. To assess the obtainability of the protein molecule in a solvent environment, SASA was employed as a useful parameter. In this study, SASA values were calculated and depicted to determine the effect of ATP and MSM binding on the solvent availability of 3VZB, as shown in Figures 4C and 4D. The plot presents a slight elevation in the SASA values of APO upon binding with ATP and MSM. This implies that the binding of ligands may expose some interior components of the protein to the surface. The SASA values demonstrated fair balance without remarkable fluctuations all through the simulation. The PDF plot clearly shows that in the case of ATP-bound 3VZB, the SASA values increased while retaining a steady distribution. The interplay between protein flexibility and compactness is well known. Furthermore, the solidity of the protein impacts its solvent approachable surface area. The radius of gyration (Rg) was determined to comprehend the impact of inhibitors on protein solidity and the SASA values are calculated to observe their effect on the approachable surface area for solvent. Kernel density estimation plots (KDE) of SASA and Rg for all systems are displayed in Figure 5. This provides definitive proof regarding the effect of inhibitors on the conformational states of 3VZP. The KDE plot of 3VZP-APO indicates a peak Rg value of 2.05 nm with a SASA value of 170.2 Å2. Similarly, for 3VZP-ATP, the KDE plot shows a peak Rg value of 2.06 nm and SASA of 172.1 Å2. Likewise, for 3VZP-MSM, the KDE plot reveals a peak Rg value of 2.04 nm and SASA of 168.1 Å2. The maintenance of intermolecular hydrogen bonding is essential for the structural integrity of proteins. The inspection of intermolecular H-bonds is broadly used to examine the firmness of the complexes formed of the protein and protein-ligand. In this study, the formation of intramolecular H-bonds formed was computed within 3VZB and compared them with those in the ATP and MSM complex which is shown in Figure 6A. The plot reveals that the H-bonds within APO remain relatively stable before and after ATP and MSM binding. The plots overlap and exhibit almost equivalent distribution during the simulation time. These results suggest that the intramolecular hydrogen bonding in APO plays a pivotal part in estimating the structural geometry of 3VZB. The binding of ATP and MSM does not appear due to disorganized intramolecular hydrogen bonding in 3VZB. However, the PDF distribution plot presents a slight reduction in hydrogen bonds within ATP and MSM, indicating the tenancy of few intramolecular are of the 3VZB binding region by the ATP and MSM which is shown in Figure 6B. Principal component analysis 2D projection plot presents the conformation sampling of APO, ATP and MSM on PC1 and PC2. To estimate the stability of the interactions between the protein and the ligand, the formation of intramolecular hydrogen bonds takes a pivotal role. In this study, we investigated the time-dependent behavior of intramolecular hydrogen bonds between ATP and MSM and plotted the results. The plot revealed that despite higher fluctuations, up to four hydrogen bonds were established between ATP and MSM. Our analysis indicates that the docked complex remained stable during the simulation, maintained by at least 3 to 12 hydrogen bonds with ATP and 1 to 3 with MSM The PDF distribution plot showed the fluctuations in the intramolecular hydrogen bonds between MSM and ATP, but they retained their stability. To explore the collective movements in APO and ATP and MSM complex, we conducted PCA. The initial eigenvectors (EVs) take a crucial role in the global motion of a protein molecule. Hence, we carried out PCA to study the conformational dynamics of APO and ATP and MSM complex during the simulation as shown in Figure 7. The time evolution of PCA suggests that the general adaptability of the ATP and MSM complex was diminished on both EVs, implying firmness in Figure 4E. The plot clearly demonstrates that the APO and ATP and MSM complex engaged almost all the conformational motions and imbricated. In general, the reduced number of movements detected in the ATP complex suggests that ATP did not significantly affect the 3VZB conformation and dynamics, thus supporting the firmness of the complex. The analysis of free energy landscapes (FELs) is a common approach to investigating protein folding mechanisms and overall stability. The FEL plots offer a visual representation of the most secure conformational arrangements within a protein structure. In this study, we produced FEL plots for PC1 and PC2. Deeper blue areas signify a more stable protein configuration with lessened energy levels.

**Figure 3.**
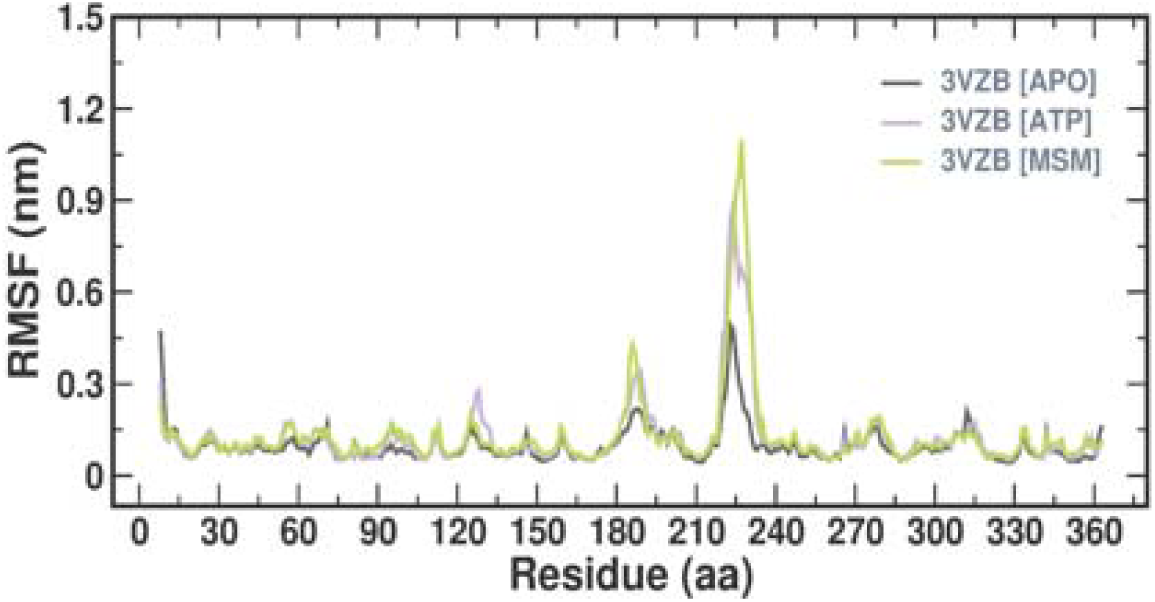
Time evolution of the RMSF values

**Figure 4.**
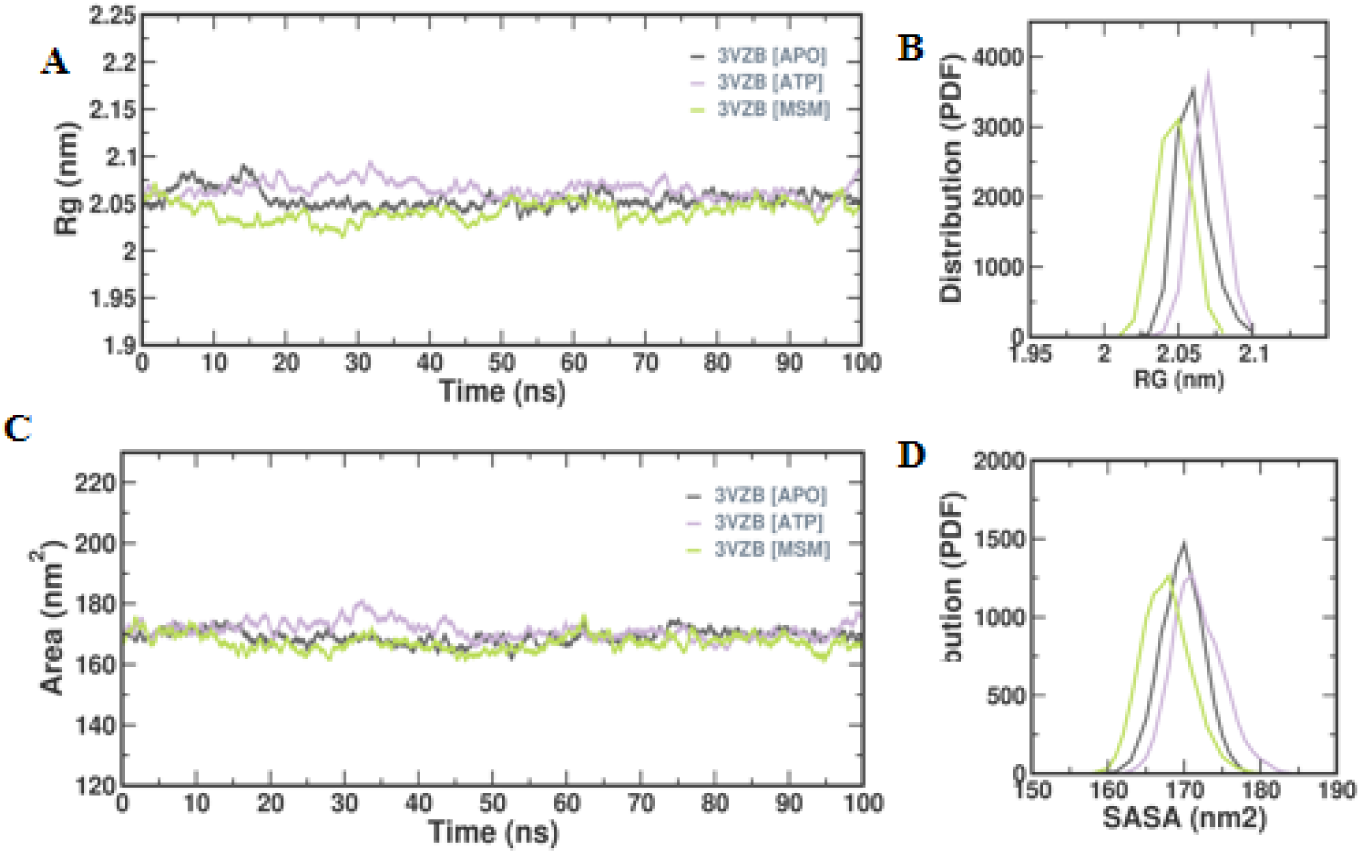
Conformational dynamics of 3VZB-APO, 3VZB-ATP, and 3VZB-MSM complex (A) Time evolution of the Rg values (B) Probability distribution of Rg values (C)Time evolution of the SASA values (D) Probability distribution of SASA values

**Figure 5.**
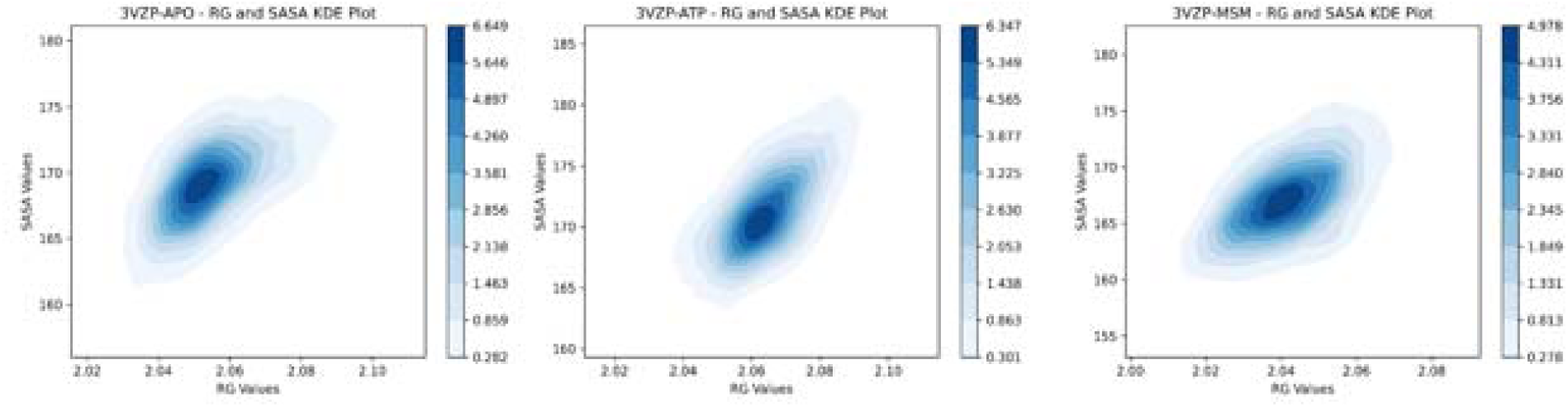
KDE Plot for RG and SASA value

**Figure 6.**
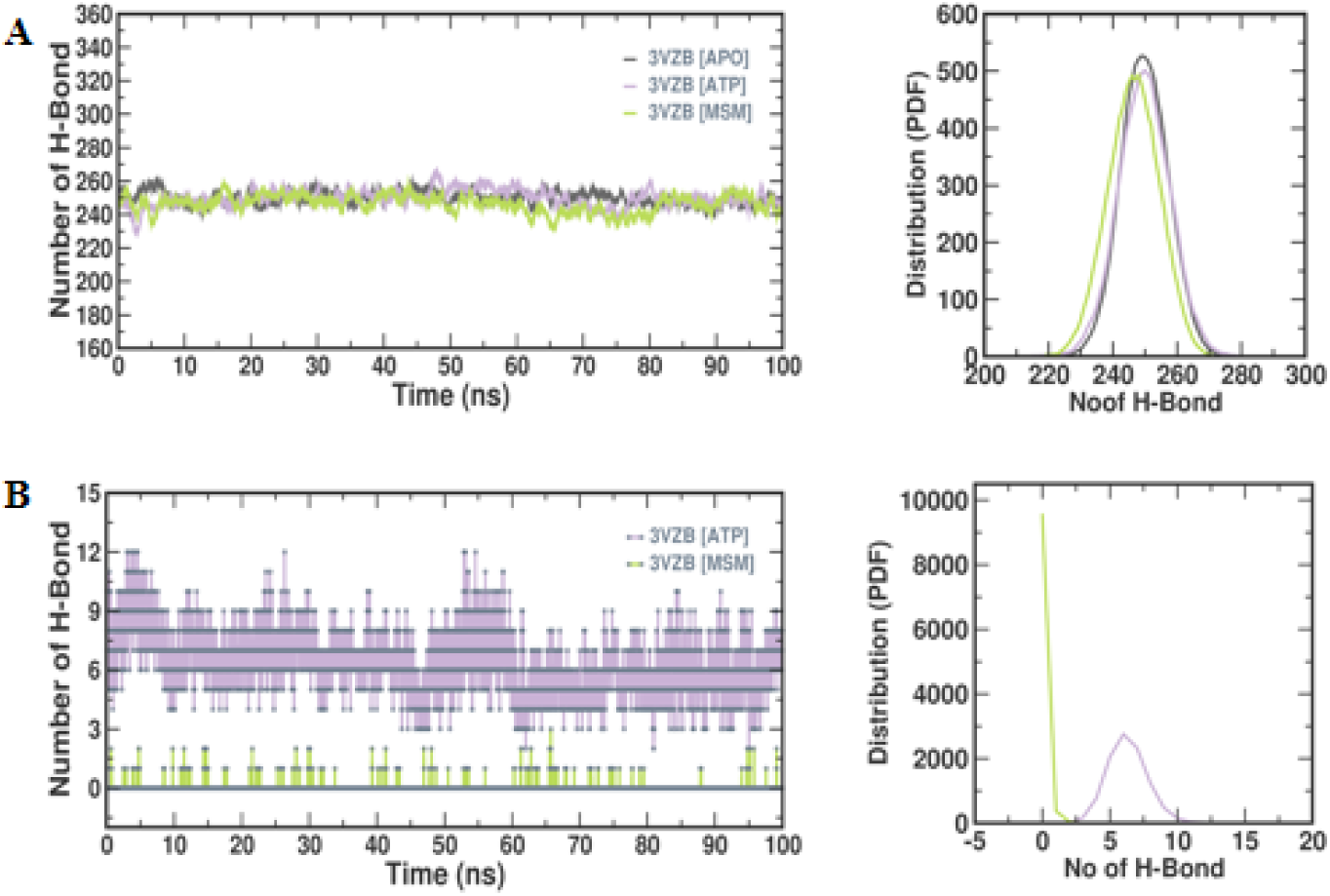
(A) Intermolecular hydrogen bonds between ATP and MSM during the simulation time and probability distribution plot of Intermolecular hydrogen bonds (B) Intramolecular hydroge hydrogen bonds between ATP and MSM during the simulation time and probability distribution plot of Intramolecular hydrogen bonds

**Figure 7.**
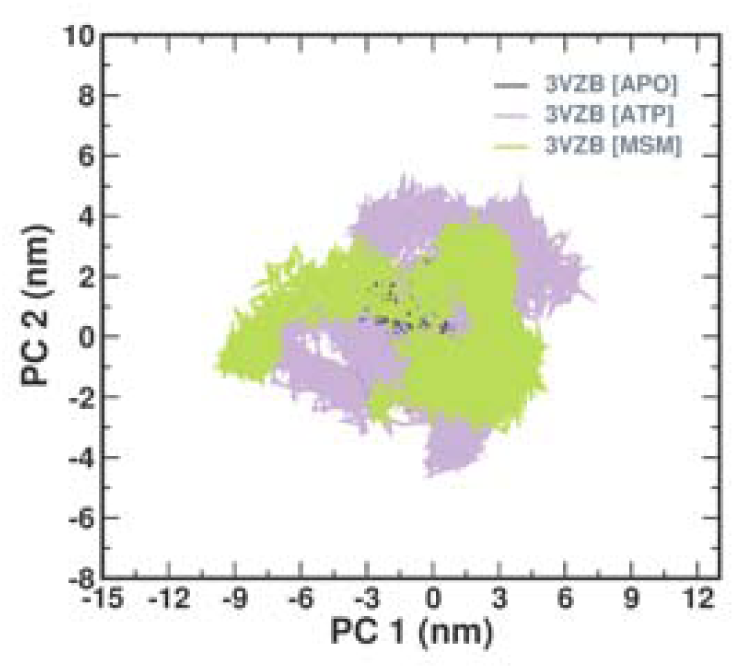
Principal component analysis 2D projection plot shows the conformation sampling of APO, ATP and MSM on PC1 and PC2

The plots indicate energy merits ranging from 0 to 16 kJ/mol throughout the simulation of APO, ATP, and MSM. The FEL plots reveal that APO, ATP and MSM complex display a single global minimum, confined to a huge local basin. These findings suggest that ATP and MSM do not cause any significant conformational changes in the 3VZB structure, thus stabilizing it which is shown in Figures 8 A, B and C. The time-evolution of secondary structure elements in the target during the protein MD is then analyzed thoroughly.

**Figure 8.**
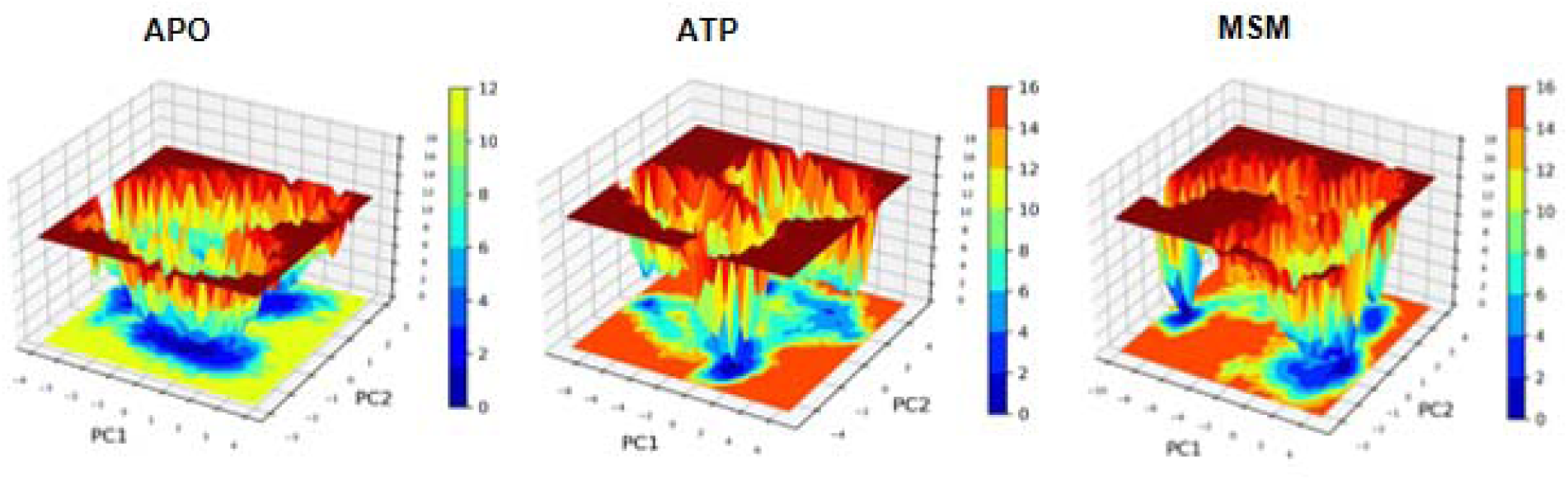
Free Energy Landscape for APO, ATP and MSM

The DSSP-annotated SSEs of the target protein 3VZPAPO and its complex with ATP and MSM systems are plotted to analyze the SSEs during the MD trajectory frames as presented in Supplementary Figure 2 and RMSD of ligand (ATP and MSM) shown in supplementary Figure 1. The important amino acid SER185 was selected for distance calculation based on the analysis the ATP had min 4 to 5Å distance, this indicates that ATP is very near to SER185. The SER185 had a distance of 40Å in the case of MSM, this indicates that MSM is moved from the active site of SER185 which is shown in Supplementary Figure 3. Concerning the estimation of the binding affinity of 3VZB-ATP and 3VZB-MSM, we inspected the relative binding power within the protein of summary energy. The comparison between the binding power of 23VZB-ATP and 3VZB-MSM regarding inhibitors calculated by the MMPBSA method and it is presented in Table 3. Over a stable simulation trajectory, the residue-level benefactions to the interaction energy are calculated. The results show that 3VZB-ATP has a van der Waals energy of -103.803 +/- 27.078 kJ/mol, an electrostatic energy of -573.197 +/- 208.560 kJ/mol, a polar solvation energy of 663.390 +/- 260.790 kJ/mol, and a binding energy of -28.730 +/- 121.725 kJ/mol. 3VZB- MSM has a van der Waals energy of -4.280 +/-7.223 kJ/mol, an electrostatic energy of - 1.741 +/- 12.220 kJ/mol, a polar solvation energy of -6.892 +/- 27.126 kJ/mol, and a binding energy of -13.922 +/- 19.518 kJ/mol.

**Table 3.**
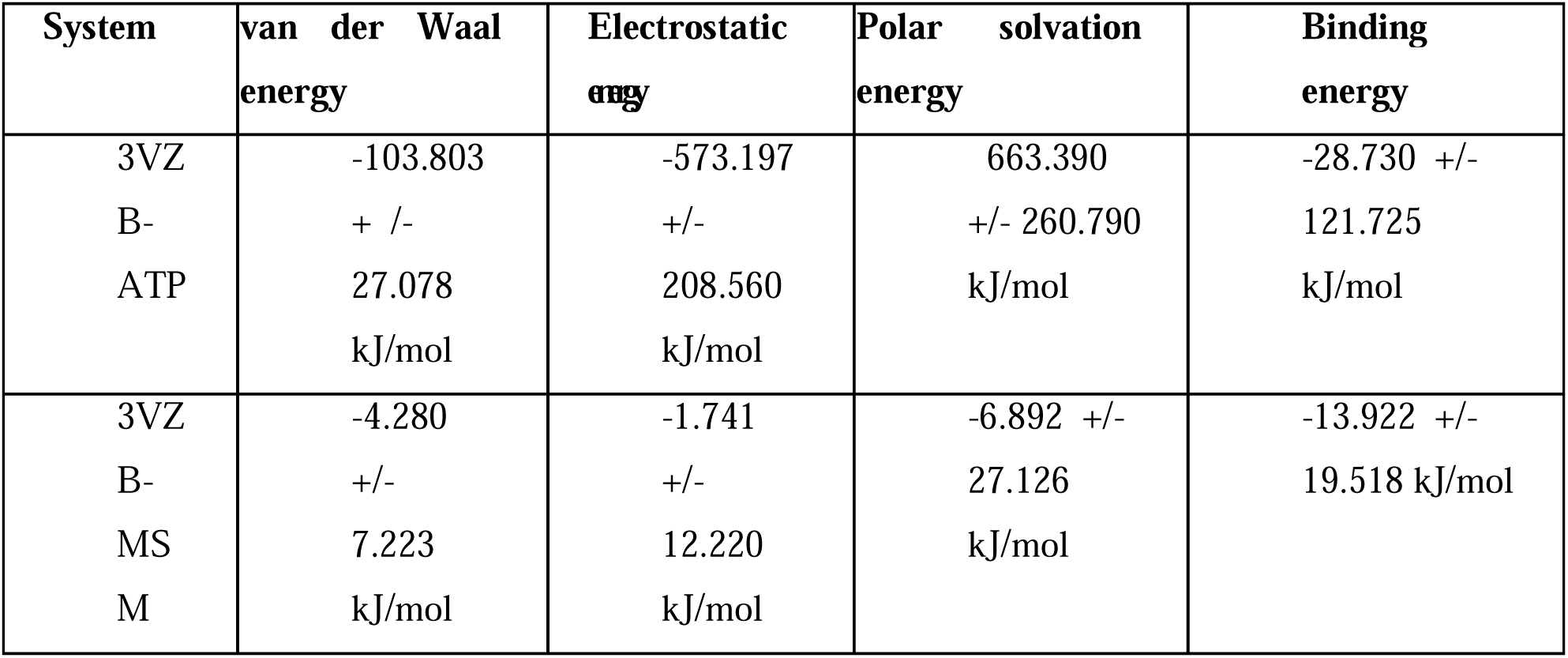
Comparison of the binding strength of 23VZB-ATP and 3VZB-MSM concerning inhibitors computed via the MM-PBSA method.

### 3.4. MSM Decreased Cell Viability and Induced Cytotoxic Effect in C6 glioma cell line

The MTT assay was used to determine MSM’s cytotoxic capability on the C6 glioma cell line. MSM is a dietary supplement and is currently prescribed for rheumatoid arthritis and various inflammatory disorders and can be used for 3 grams/day [4]. In our study, C6 glioma cells were displayed to different concentrations of MSM, which range from 2.5 to 2000 mM/mL, as at high doses, MSM is not highly toxic in normal cells. MSM prevented the proliferation of C6 glioma cells in a concentration dependent manner with minimum inhibitory concentrations (IC50) value of 393.459 mM/mL in C6 cells which is shown in Table 4. Thus, the results presented that MSM flaunted significant cytotoxicity in C6 glioma cells only at elevated densities (600 mM/mL). Previous findings proved that MSM has anti-cancer properties with no adverse effect on gingival squamous adenocarcinoma and 200 mM MSM prevents cell viability in a concentration dependent manner on gingival cancer cell line YD-38[22]. In another study, Eun et al. observed that MSM suppresses breast cancer growth in Bal b/c athymic nude mice inoculated with MDA-MB 231 cells by decreasing the expression of STAT3, STAT5b, IGF-1R and VEGF. In this study, MSM was used at 300 mM and 500 mM against human breast cancer cell lines MDA-MB 231 and SK-BR3 and thus, MSM is effective in breast cancer too [23]. MSM in combination with a low concentration of allicin, can be used for breast cancer treatment. All the previous findings support that MSM is a potential dietary supplement which can be used for the treatment of various cancer even GBM as well as it is not toxic at high doze.

**Table 4:**
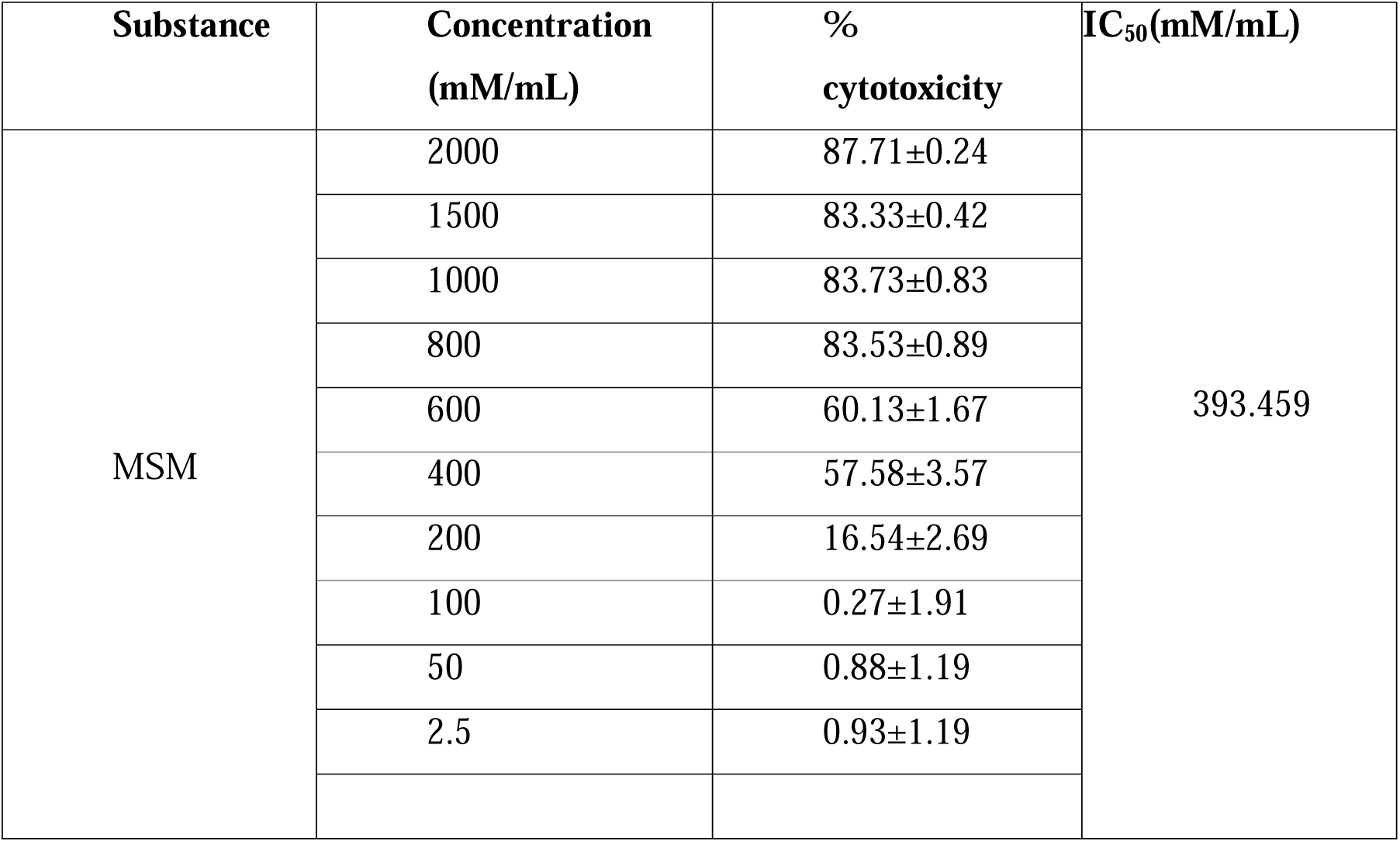
Analysis of the Inhibitory properties of MSM against the C6 cell line by MTT assay.

| Substance | Concentration<br>(mM/mL) | %<br>cytotoxicity | IC<sub>50</sub>(mM/mL) |
| --- | --- | --- | --- |
| MSM | 2000 | 87.71±0.24 | 393.459 |
|  | 1500 | 83.33±0.42 |  |
|  | 1000 | 83.73±0.83 |  |
|  | 800 | 83.53±0.89 |  |
|  | 600 | 60.13±1.67 |  |
|  | 400 | 57.58±3.57 |  |
|  | 200 | 16.54±2.69 |  |
|  | 100 | 0.27±1.91 |  |
|  | 50 | 0.88±1.19 |  |
|  | 2.5 | 0.93±1.19 |  |

### 3.5 MSM Induced Morphological Changes in C6 glioma cell line

The cellular morphology of C6 glioma cell line upon treatment of 400 and 600 mM/mL of MSM was examined employing an inverted phase contrast microscope. Phase contrast photomicrographs clearly portrayed the morphological changes in the treated cells as compared to the control. When the C6 glioma cell line was treated with MSM at 400 mM, we observed morphological changes in cells, i.e. reduction in the number of pleomorphic cells, as well as nuclei size is reduced and in the case of 600 mM of MSM, the nuclei seemed very small in size which is visible as light dark brown colour and reduction in the number of pleomorphic cells and entire morphology changed in comparison to untreated C6 glioma cell line which is shown in Supplementary Figure 4 A, B and C for different treatment conditions with and without MSM. As an outcome, we chose the best concentrations depending on the results above of MSM (400 and 600 mM/mL) to carry on with our study.

### 3.6 MSM Induced DNA Fragmentation in C6 glioma cell line

One of the characteristics of apoptosis is DNA fragmentation; consequently, we proceeded with a DNA fragmentation ladder assay to detect whether MSM treatment induces DNA fragmentation in C6 glioma cell line. MSM induced apoptosis, and DNA was degraded by DNA activated DNase (CAD). In this study 1kb marker is on the left, and the 100bp marker is on the extreme right, whereas the control DNA on the right and in between our C6 cell line treated with MSM at 400 mM and 600 mM is visible on the DNA. When we compared the result with previous findings, we noted that MSM was used in combination with tamoxifen which inhibited tumour growth and metastasis. MSM at 200 mM had the potential to induce apoptosis compared to individual agents which were confirmed by DNA fragmentation assay[22] as well as in another study conducted by Madhuri et al shown that Bacoside A treated U87 MG cells show DNA ladder formation[24] whereas, in this study, we observed that the spread out dark stained portion increases with a rise in MSM concentration as compared to the control which is clearly visible. This proves that MSM inhibits apoptosis at 400 mM and 600 mM concentrations respectively and is shown in Figure 9 as DNA strand fragmentation are the hallmark of apoptotic cell death.

**Figure 9.**
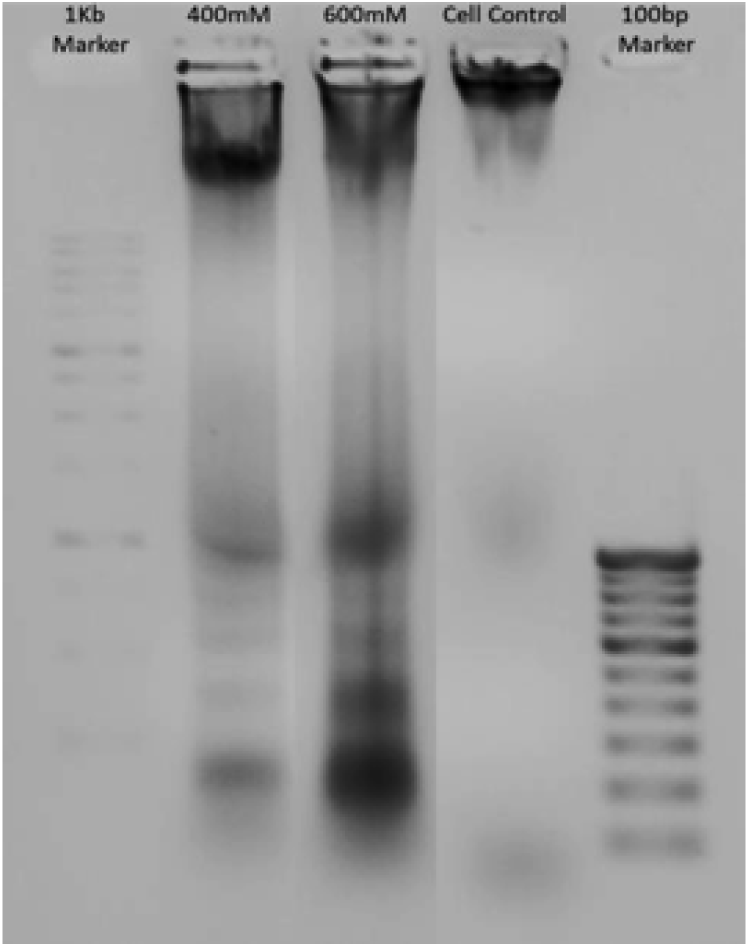
Detection of DNA Fragmentation of MSM in C6 glioma cell line

### 3.7 MSM Augmented Intracellular ROS Generation in C6 glioma cell line

In order to support our study, we looked further into how MSM affected the intracellular redox status in C6 glioma cell line. According to studies, the beginning of apoptosis is related to augmented intracellular ROS production [1]. We performed Dichlorodihydrofluorescein diacetate (DCFHDA) staining and detected a fluorescence intensity to produce ROS in the C6 glioma cell line which showed that in control, production of ROS showed a fluorescence intensity of 54.765 a.u. whereas, the production of ROS at 600 mM was 128.335 a.u. The ROS activity at 400 mM showed a fluorescence intensity of 76.255 au. which showed many active cells showing green fluorescence indicative of slightly lesser ROS activity. Thus, fluorescent micrographs depict that MSM enhanced ROS activity in C6 glioma cell line in a concentration (dose)-dependent manner which is shown in Figure 10 A, B and C and graphical representation of the quantified data is presented in Figure 10i. Previous findings show that in 2016 for the first time, Arzu et al observed that MSM induced apoptosis in HCT116 colon cancer cells [26]. Dong et al. also observed that MSM regulates the activation of STAT5b and decreases HER2 expression [27] even MSM induces apoptosis in liver cancer [28]. MSM is known to decrease the viability of prostate cancer cells and the arrest of the cell cycle progression in the G0/G1 phase through the induction of apoptosis. MSM at a dose of 200 mM decreases the viability and invasiveness of prostate cancer cells and there is a possibility that MSM can be used for the treatment of prostate cancer in future [29]. These findings prove that elevated production of ROS in the glioma cultures in vitro is the hallmark of apoptosis and can be used as a positive direction towards control of brain tumor progression

**Figure 10.**
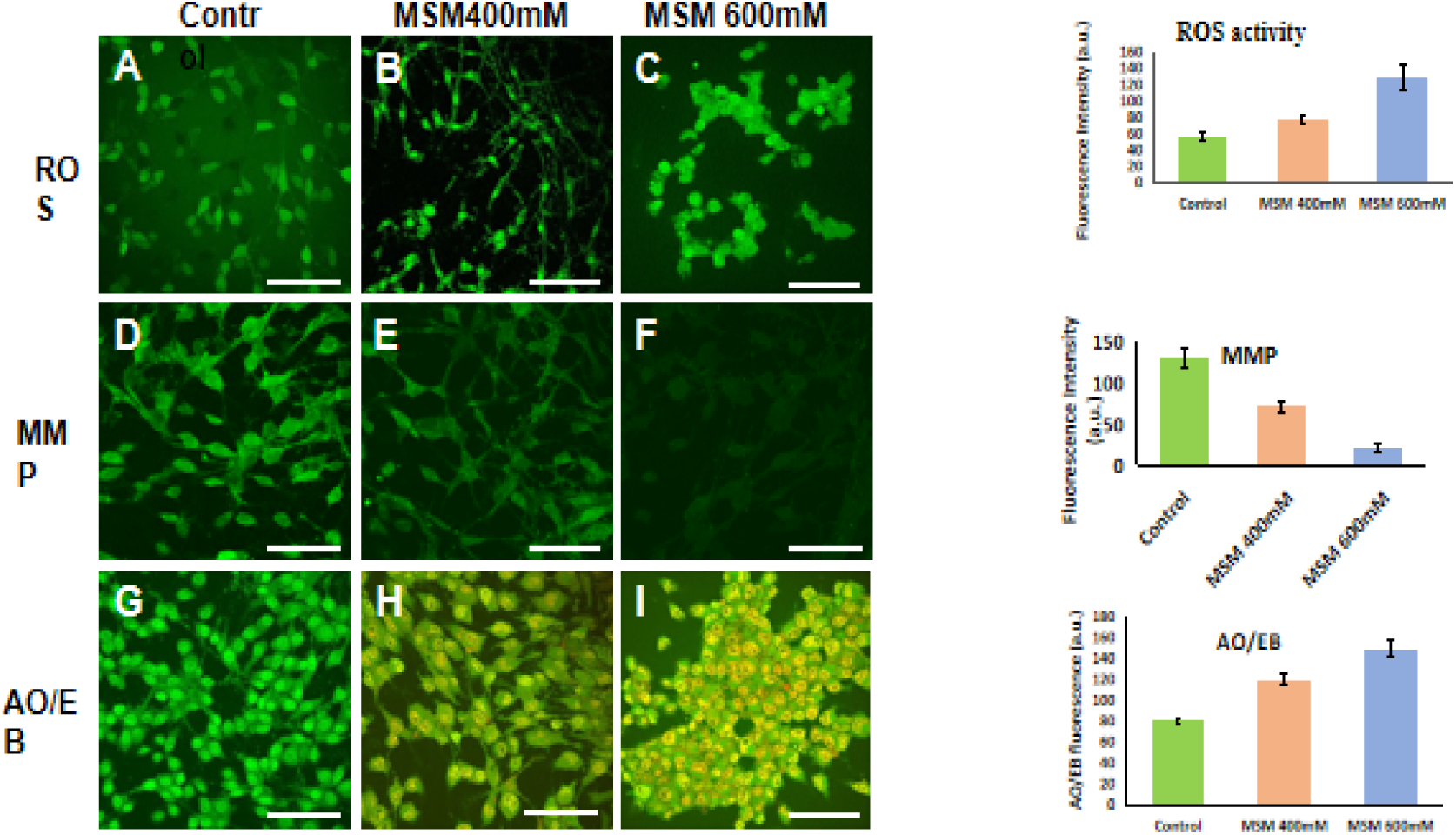
(i)Fluorescence microscopy images of the C6 glioma cell line subjected to DCFH-DA staining depicting control as A, 400 mM at B and 600 mM at C (ii) Fluorescence microscopy images of the C6 glioma cell line subjected to Rhodamine 123 dye control as D, 400 mM at E and 600 mM at F (iii) Fluorescence microscopy images of the C6 glioma cells subjected to AO/EB staining control as G, 400 mM as H and 600 mM as I

### 3.8 MSM Disrupted Mitochondrial Membrane Potential (MMP) in C6 glioma cell line

A crucial stage in the mitochondrial apoptotic cascade is the derangement of the mitochondrial membrane, which is followed by the dropping of the φ Rhodamine 123 (Rh123) is a cationic green dye and hence, localizes in mitochondria and it can be used to detect mitochondrial potential as mitochondrial dysfunction which can lead to apoptosis [1]. The decrease in Rh123 retention and fluorescence has been demonstrated to be accompanied by disruption of mitochondrial potential. In this study, the control or untreated gliomas are in extreme left-hand side as shown in Figure 10D having the highest fluorescent intensity of 129.813 a.u. which is indicative of highest mitochondrial potential. When cells were treated with MSM at 400 mM and 600 mM and stained with Rh123, we observed fluorescent dark green intensity decreases at 400 mM treated cells with a decrease in retention to 71.571 a.u. whereas, at 600 mM, the fluorescence signal from Rh123 is hardly visible with a drastic decrease of retention value to 22.961 a.u. as shown in Figure 10E and F. This proves that treating cells with MSM at 400mM and 600 mM increases the disruption of mitochondrial potential, which is shown in Figure 10 and the quantified data is shown in form of graph in Figure 10 ii. This proves that MSM decreased the φ in a dose-dependent manner and MSM disrupted MMP in C6 glioma cell line. Similar to our findings, in one of the studies by Karolina et al, they observed that MSM at 300 mM acted as a sensitizer of endometrial cancer (EC) cells to doxorubicin, for which EC cells quickly acquire resistance. DNA damage was observed and gave a positive response when MSM at 400 mM was used in combination with doxorubicin [30]. In another study, Doh et al observed that MSM diminished the viability of HT-29 cells and augmented arrest of the cell cycle progression at G0/G1 phase and apoptosis as well as vanquished sphere-forming ability and reduces the expression of stemness markers in HT29 cells [31]. Finally, based on these previous findings as well as our findings prove that mitochondrial membrane disruption is one of the important stages in mitochondrial apoptotic cascade and in our study, φ is reduced in a dose-dependent manner for the MSM treatment.

### 3.9. AO/EB staining of C6 glioma cell line with MSM

Finally, we performed AO/EB staining, in which live cells appear fluorescent green while late apoptotic cells show condensed and fragmented yellowish-orange chromatin. The findings demonstrated that a rise in MSM concentration causes a steady increase in orange and red staining, together with a reduced fluorescence green of nuclei, which denoted cell damage and apoptosis. We observed fragmented and limited numbers of yellowish-orange stains with increased fluorescent intensity of 119.129 a.u. at 400 mM MSM treated cells, while at 600 mM, we observed a large number of the yellowish-orange nuclei increasing the fluorescent intensity to 148.816 mM, which is shown in Figure 10G, H and I and the results are depicted in form of graph in Figure 10D when compared to the control untreated gliomas having no fragmented chromatins and as a result showing a lesser fluorescent intensity of 79.974 a.u. which is indicative of no nuclear damage. MSM had a better anti-cancer effect on human hepatocellular carcinoma (HepG2) in comparison to human gastric carcinoma (AGS) and human esophageal squamous cell carcinoma (KYSE-30) as IC50 of MSM on HepG2 is least with 21.87 mg/ml followed by 28.04 mg/ml and 27.98 mg/ml after 72 hr and in this study, during AO/EB staining, the results show that at a concentration of 21-29 mg/ml of MSM, the cells show morphological changes [31]. Nuclear deformation demonstrated the induction of apoptosis, showing the anti-proliferative actions of MSM against glioma cells. **Figure 10-** (i)Fluorescence microscopy images of the C6 glioma cell line subjected to DCFH-DA staining depicting control as A, 400 mM at B and 600 mM at C (ii) Fluorescence microscopy images of the C6 glioma cell line subjected to Rhodamine 123 dye control as D, 400 mM at E and 600 mM at F (iii) Fluorescence microscopy images of the C6 glioma cells subjected to AO/EB staining control as G, 400 mM as H and 600 mM as I

## 4. Future Direction and Conclusion

In this study, we proved the binding mechanism of MSM with Sphk1. MSM actively interacts with the active site residues of Sphk1 through van der Waals interactions and MSM controls tumour growth and exhibits tumoricidal effects in the C6 glioma cell line at 600 mM. For the first time, MSM is investigated against C6 glioma cell lines, but still in-depth in vitro and in vivo studies need to be conducted before we proceed with the clinical trials. This study supports that MSM as well as other anti-inflammatory dietary supplements can be used not only for GBM but for different types of cancer and neurodegenerative diseases as even at high dose dietary supplements are not toxic and will not cause adverse effect which can cause severe damage to body. MSM can act as a promising Sphk1 inhibitor which is also involved in GBM as well controls tumor growth in C6 glioma cell line. This study opens a door for researchers to explore other dietary supplements for different fatal disease.

## Abbreviations

Sphk1: Sphingosine kinase 1
MSM: Methylsulfonylmethane
GBM: Glioblastoma Multiforme
FDA: Food and Drug Administration
VEGF: Anti-vascular endothelial growth factors
TTF: tumour treating field therapy
HCC: Hepatocellular carcinoma
AOEB: Acridine orange and Ethidium Bromide
ROS: Reactive Oxygen Species
ACPYPE: AnteChamber PYthon Parser interface
DCFH –DA: Dichlorodihydrofluorescein diacetate
MMP: Mitochondrial membrane potential
MD: Molecular Dynamics
SASA: Solvent accessible surface area
KDE: Kernel density estimation
RMSD: Root mean square deviation
RMSF: Root mean square fluctuation
PCA: Principal Component Analysis
Rg: Radius of Gyration
BBB: Blood Brain Barrier
Rhodamine 123: Rh123
CAD: DNA-activated DNase

## Data availability statement

All data that support the findings of this study are included within the article (and any supplementary files).

## Acknowledgement

We would like to thank the Radiant Research for providing us in vitro facilities

## Funding

Self-funded

## Authorship contribution statement

Faizan Ahmad – Conceptualization, in silico and in vitro aanalysis, writing and editing, Anik Karan – conceptualization, supervising, reviewing and editing, Richard Jayaraj - supervising, reviewing and editing

## Ethical Approval

Not applicable

## Consent to Participate

Not applicable

## Consent for Publication

All authors have given consent for publication

## Competing Interests

The authors declare they have no conflict of interest

## Graphical Abstract

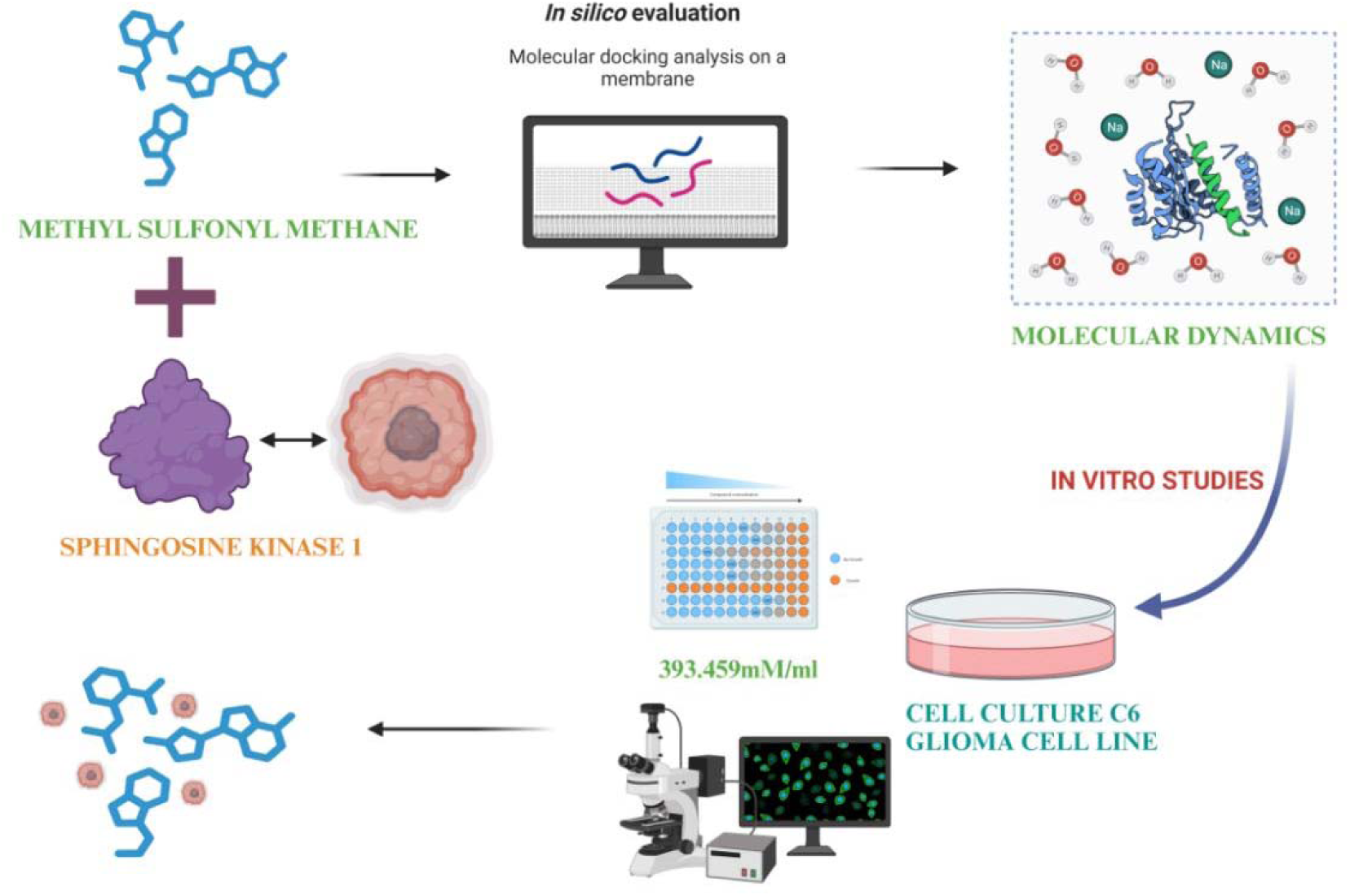

## Supplementary Figures

**Figure S1.**
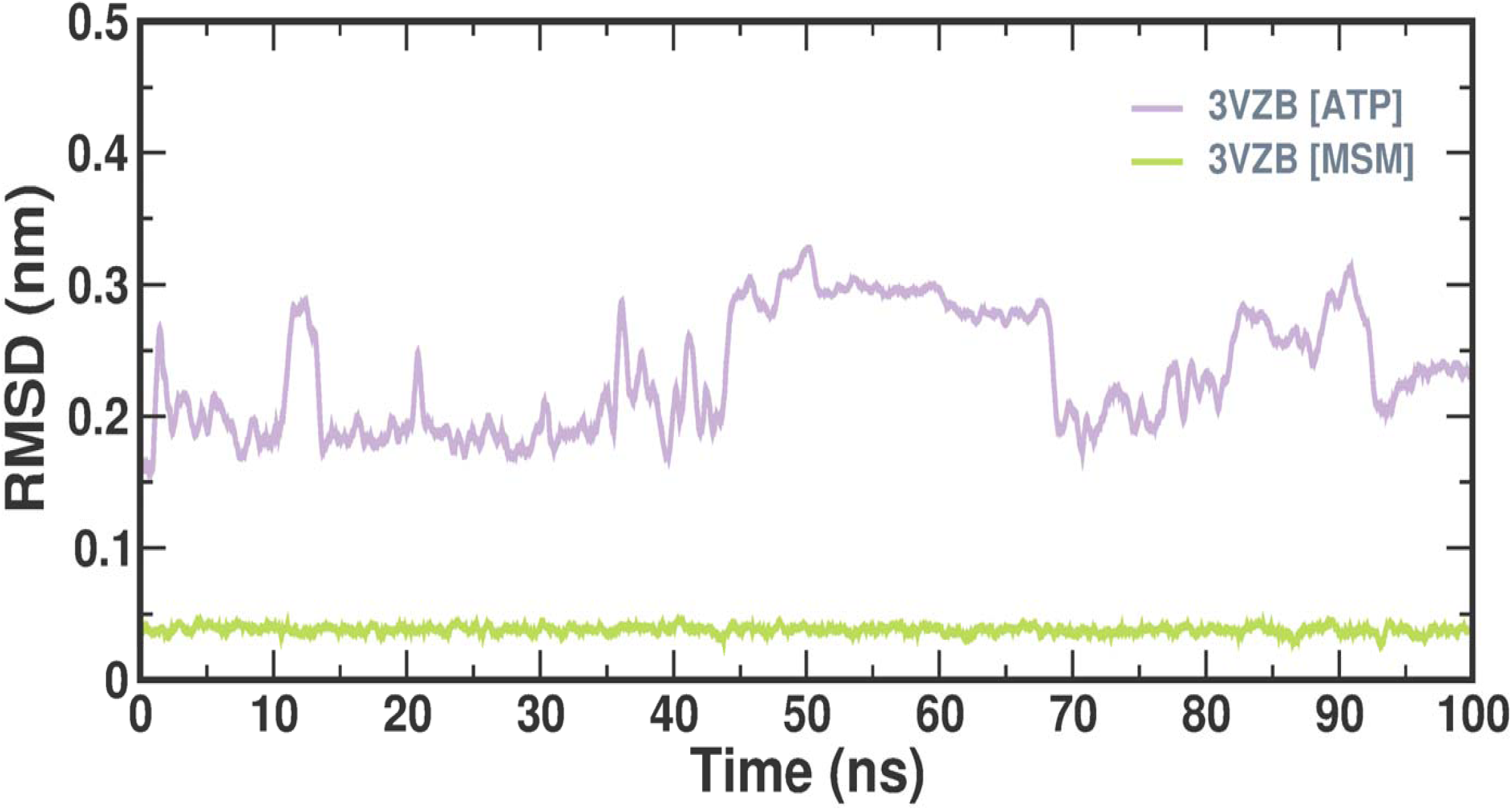

**Figure S2.**
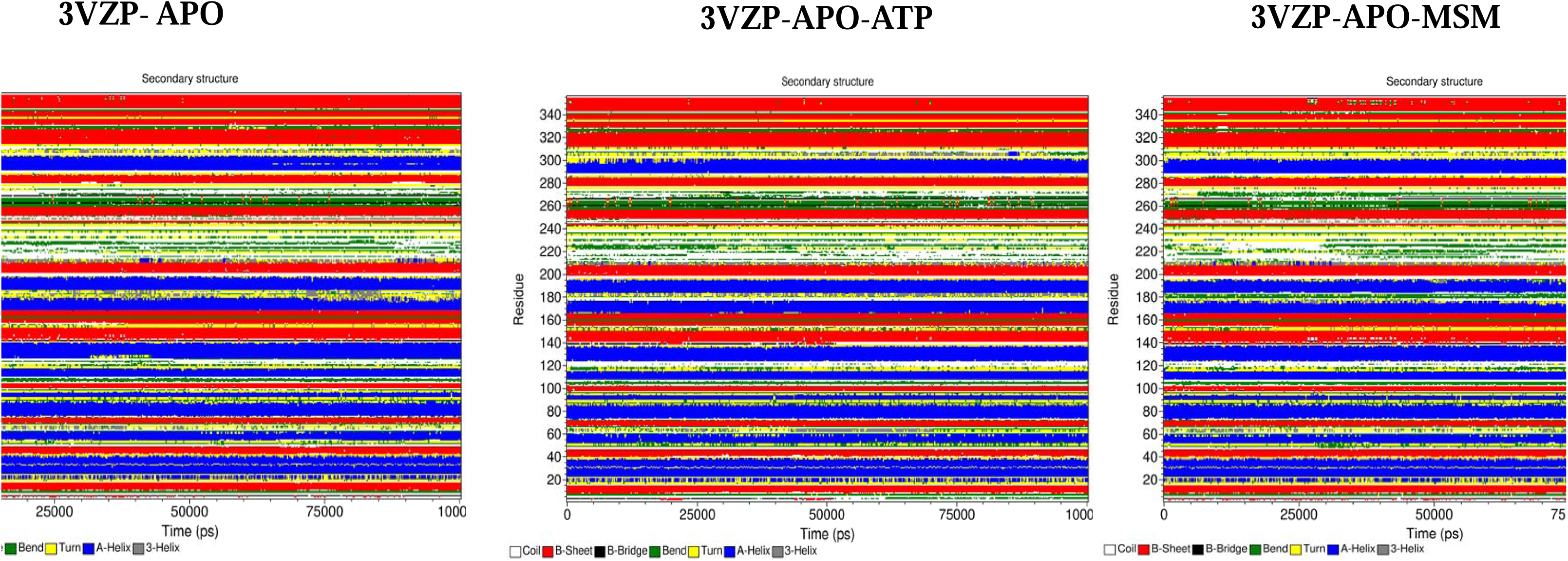

**Figure S3.**
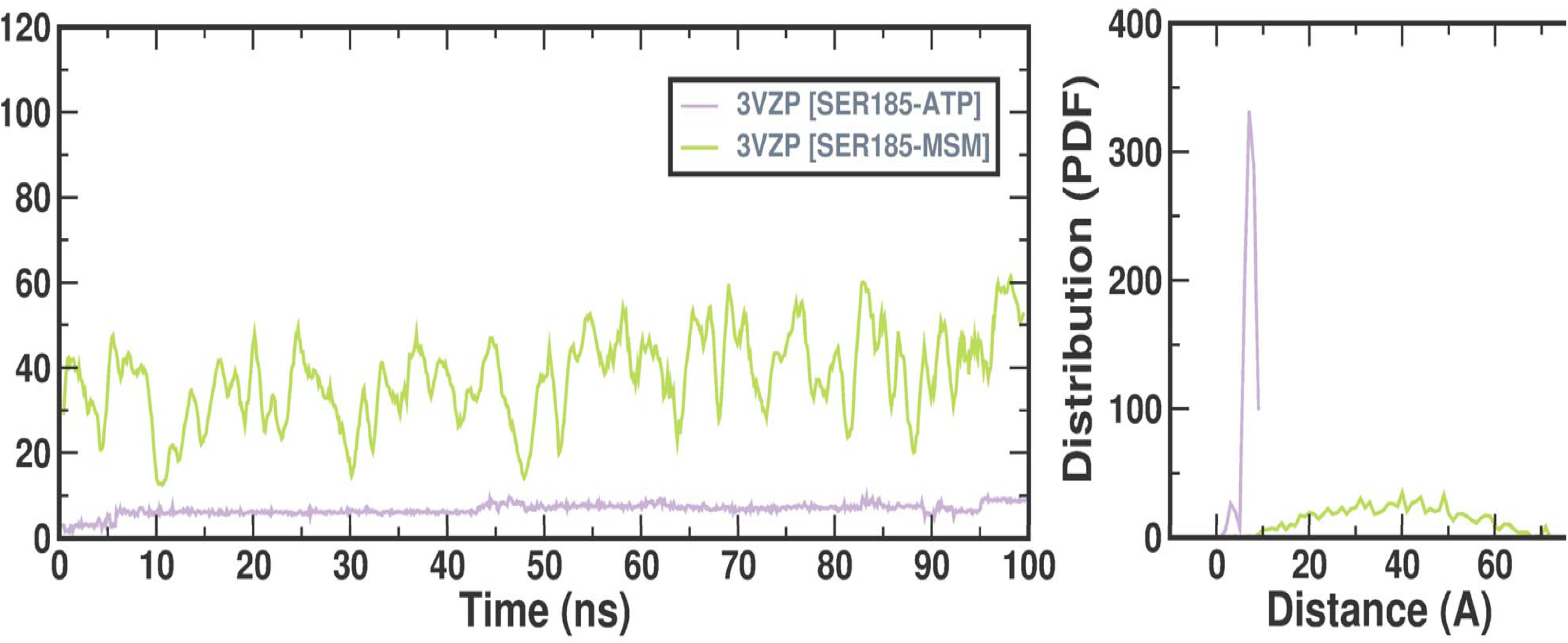

**Figure S4.**
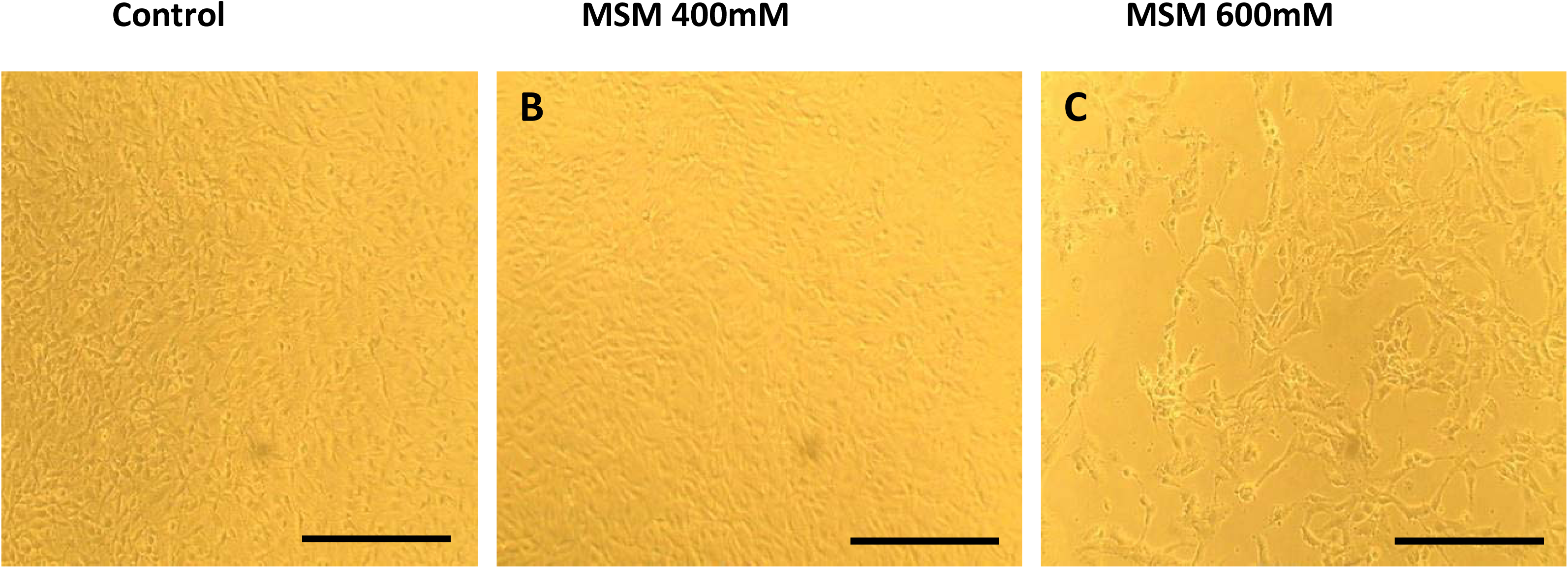

## Notes

### Competing Interest Statement

The authors have declared no competing interest.

