## Supplementary 1 for "Methylsulfonylmethane: A Potential Dietary Supplement targeting sphingosine kinase 1 involved in Glioblastomamultiforme"

**Keywords:** SphK1, Methylsulfonylmethane (MSM), Glioblastoma multiforme, Dietary supplement, Therapeutics

**SUPPLEMENTARY DATA AND FIGURES**

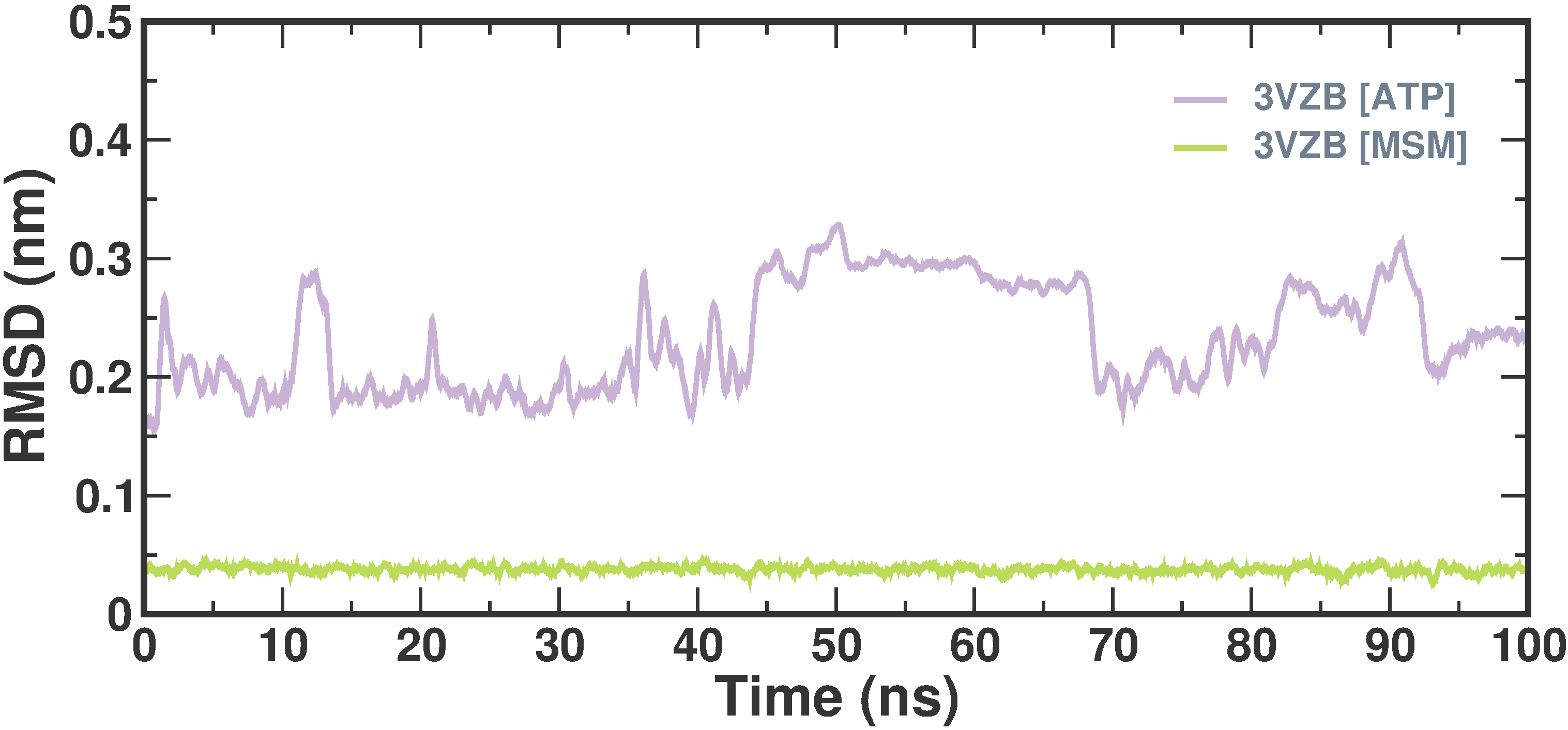

**Supplementary Figure 1**- RMSD of Ligand (ATP and MSM)

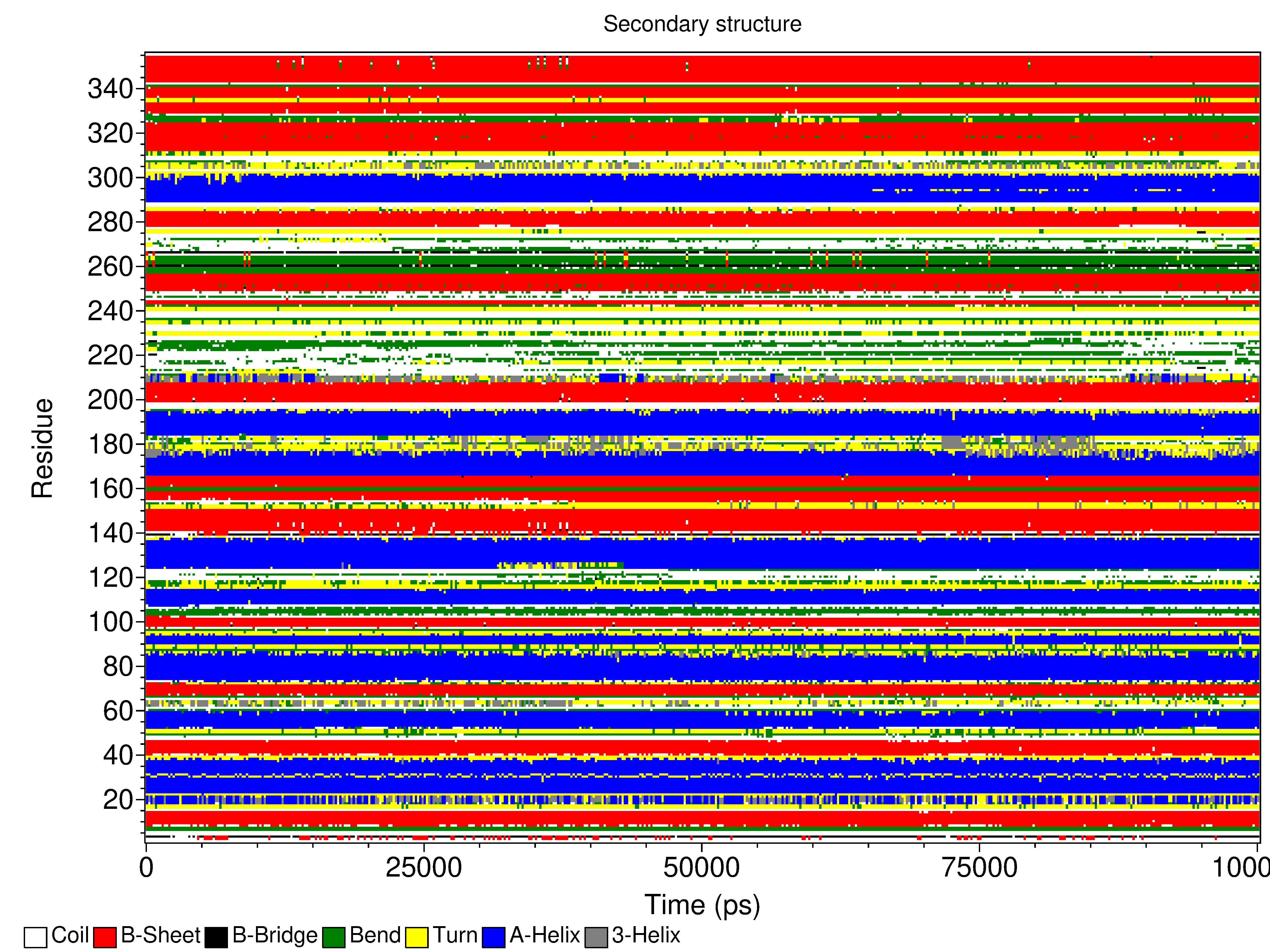

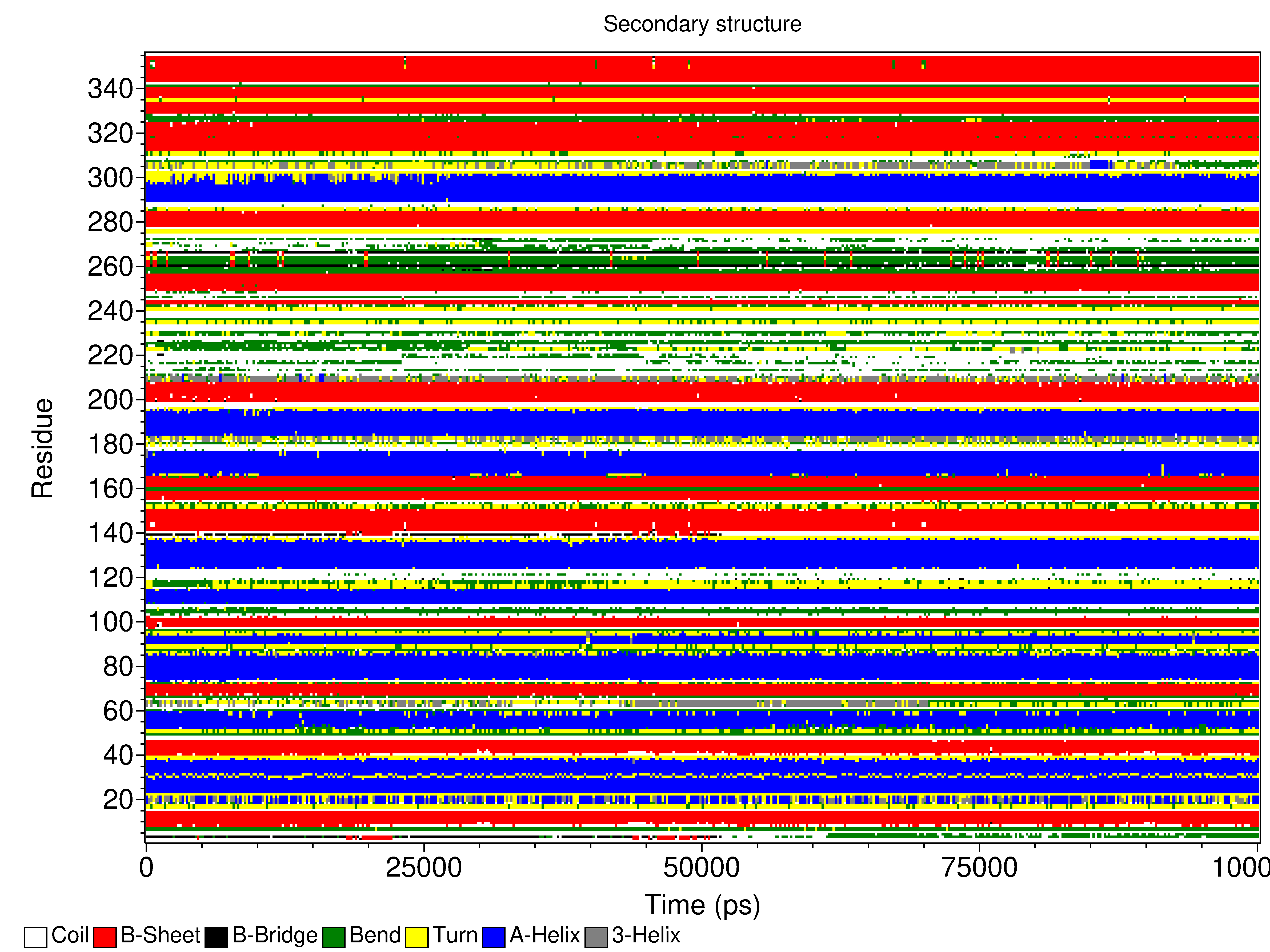

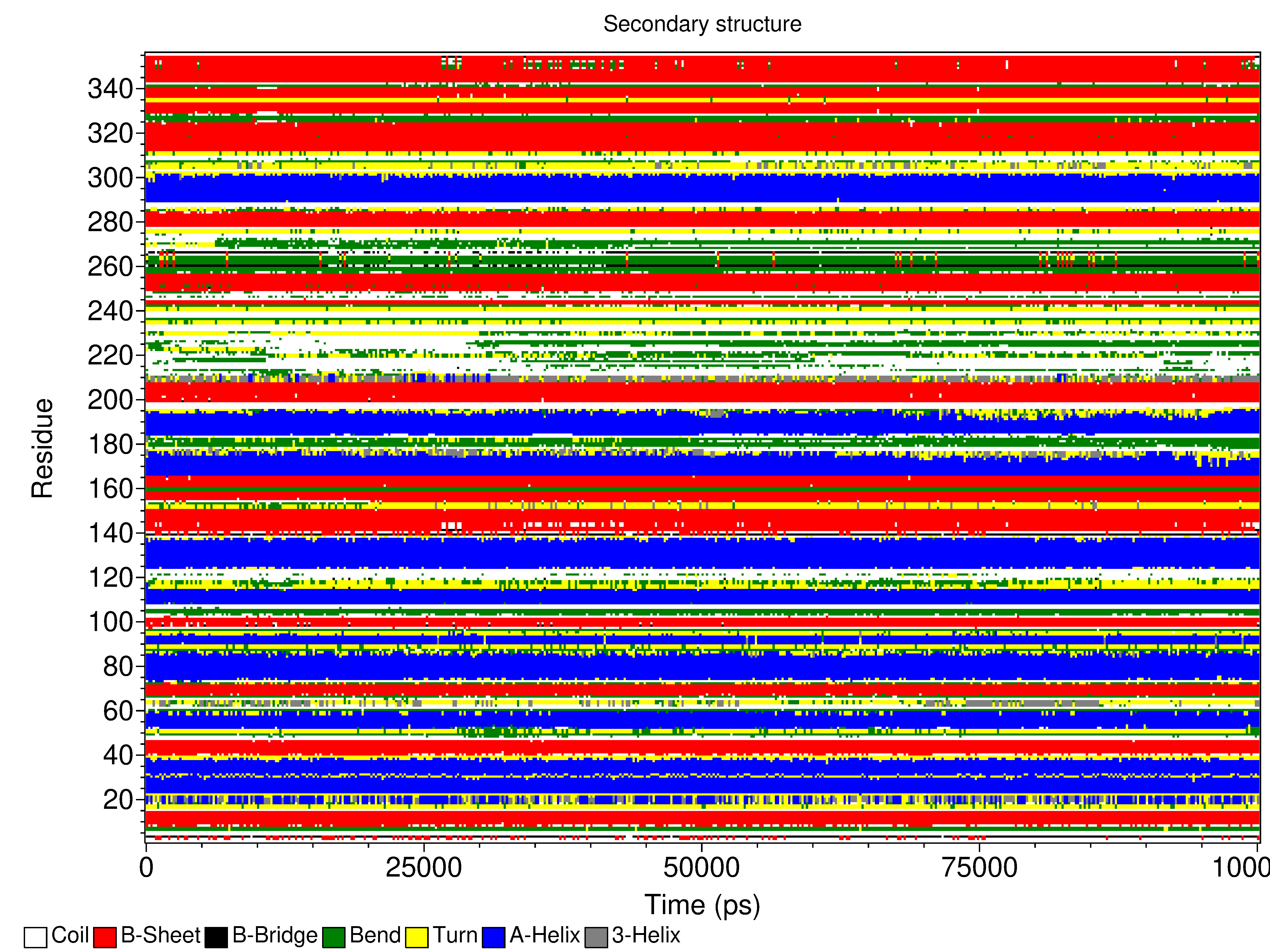

**Supplementary Figure 2-** SSEs of the target protein 3VZP-APO and its complex with ATP and MSM

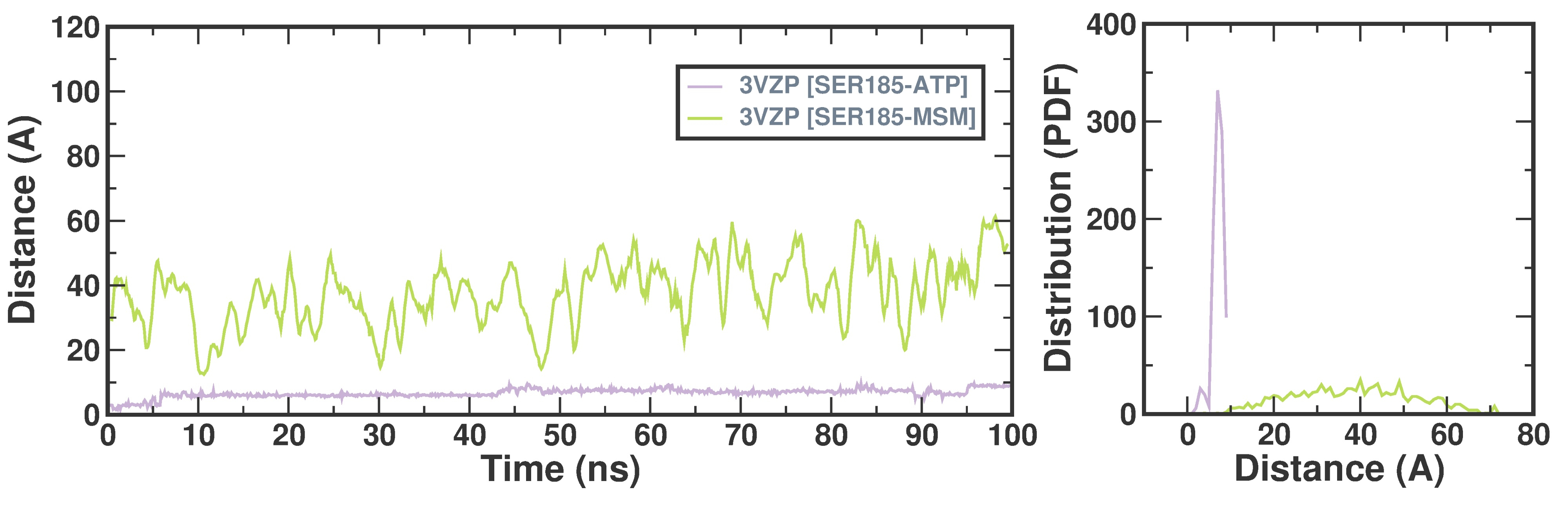

**Supplementary Figure 3 -** Distance plot for active site residue SER185 with ATP and MSM

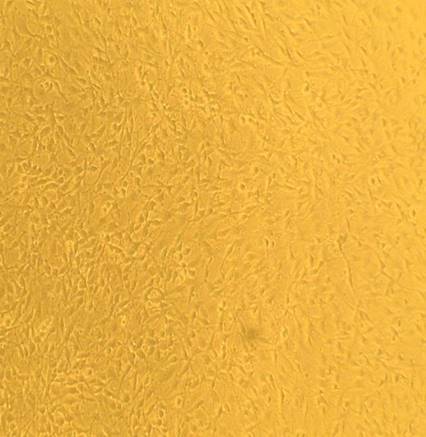

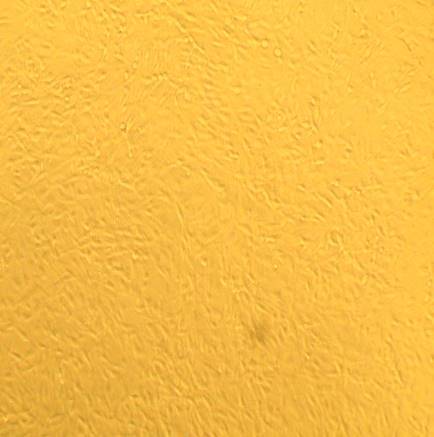

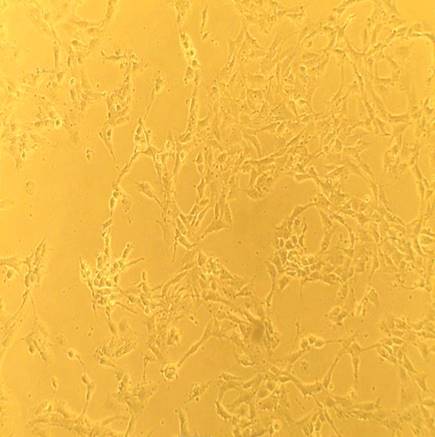

**Control**

**MSM 400mM**

**MSM 600mM**

**A**

**B**

**C**

**Supplementary Figure 4-** Phase contrast images of C6 glioma cells treated with MSM
